# Biophysical basis for the induction of glioblastoma-like phenotype in astrocytes

**DOI:** 10.1101/2023.07.31.551259

**Authors:** Laurent Starck, Melika Sarem, Bernd Heimrich, Ritwick Sawarkar, Marie-Françoise Ritz, Gregor Hutter, V. Prasad Shastri

**Affiliations:** Institute for Macromolecular Chemistry, University of Freiburg, 79104 Freiburg, Germany; BIOSS Centre for Biological Signalling Studies, University of Freiburg, 79104 Freiburg, Germany; Faculty of Medicine, Department of Neuroanatomy, University of Freiburg, 79104 Freiburg, Germany; Institute for Genetics and Medical Research Council Toxicology Unit, University of Cambridge, Cambridge, CB2 1QR United Kingdom; Brain Tumor Immunotherapy Lab, Department of Biomedicine, University of Basel, Basel, Switzerland; Department of Neurosurgery, University Hospital Basel, Basel, Switzerland

## Abstract

While the direct biological factors underlying the progression of GBM, an aggressive form of brain cancer, have been extensively studied, emerging evidence suggests that indirect biological triggers, such as traumatic brain injury (TBI), may also have a role. Since reactive astrocytes are associated with TBI, and astroglial cells are the source of proteoglycans which contribute to changes in biophysical characteristics (stochastic topography, stiffness) of the brain, we postulated a role for stochastic nanoroughness in the induction of glioma. Using a model system to emulate such physical cues, we demonstrate that human cortical astrocytes undergo spontaneous organization into spheroids in response to nanoroughness and retain the spheroid phenotype even upon withdrawal of the physical cues. Furthermore, spheroids serve as aggregation foci for naïve astrocytes; express activated MMP2, and disseminate upon implantation in mouse brain. RNA-seq revealed a tumoral phenotype with a gene expression pattern involving p53, ADAMTS proteases and fibronectin. Moreover, nanoroughness mediates a cross-talk between cancer cells and astrocytes through induced senescence. These findings implicate a role for stochastic biophysical cues in driving a potential malignant transformation of astrocytes.

## Introduction

Tumors of the CNS, accounting for approximately 2% of the neurological diseases and less than 2% of diagnosed cancers, have become a significant public health concern with a marked 17% increase in incidence since 1990 ^1^. GBM characterized by its aggressiveness, limited treatment options, and low life expectancy presents a formidable challenge in the field of neuro-oncology ^2^. While numerous factors contribute to the progression of GBM, including soluble signals and alterations in astrocyte phenotype, recent studies have shed light on the potential role of indirect biological triggers. Notably, it has been suggested that traumatic brain injury (TBI) increases the predisposition to gliomas later in life^3, 4^. While one noteworthy study has concluded an absence of such a correlation between TBI and primary brain cancer^5^, reports have also emerged describing GBM formation directly at the site of injury, years following the traumatic event^6, 7^, and radiological evidence from several case reports supports this correlation between TBI and glioma^6, 8^. Moreover, pediatric brain trauma has been associated with a twofold increase in the likelihood of brain cancer development^9^. Given the strong association between cancer and inflammation^10^, it is plausible that trauma-induced inflammation may promote progenitor cell transformation, potentially contributing to glioma initiation^3, 11^. Immunohistochemical studies have revealed reactive astrocytes (tumor-associated astrocytes, TAAs) surrounding glioblastoma, which interact closely and communicate with cancer cells by producing proteases, cytokines, and growth factors. Initially, astrocyte reactivity appears to restrict the glioma progression; however, it ultimately favors invasion in the later stages of the disease^12^, suggesting a dual role of astrocytes, in initiating and maintaining gliomas^13^.

Mechanobiology, the interplay between physical input and biological processes, represents a prominent alternative mechanism for modulating cell behavior^14^. Over the past two decades, a clear picture of the role of mechanobiology in cell phenotype and fate has emerged^15–18^, and changes to matrix stiffness has been linked to cancer progression^19–21^. Despite the brain’s characteristic softness (E modulus 1.3 – 1.9 KPa)^22, 23^ and the absence of substantial fibrous extracellular matrix proteins, brain tissue is rich in chondroitin sulfate and heparin sulfate proteoglycans (PGs) secreted by astrocytes. Proteoglycans can form large aggregates with hyaluronic acid, contribute significantly to the microstructure of the brain^24^ and present nanoscale physical cues in the form of stochastic nanoroughness. Previous studies have demonstrated the impact of stochastic nanoroughness on stem cell differentiation, morphology, and extracellular matrix assembly^25, 26^. Notably we have shown that stochastic nanoroughness impacts the lineage commitment of telencephalic neural stem cells, and additionally, mediates and alters hippocampal neuron and astrocyte behavior via the mechanosensing channel protein Piezo-1. Furthermore, our investigations have also revealed altered nanoroughness in Alzheimer’s plaques, suggesting a potential role for tissue topography in degenerative processes^27^. Blunt insult to the brain often leads to swelling^28^ and increased expression of PGs by reactive astrocytes ^29^ resulting in alterations to the physical space^30^ and can impact the mechanical properties of the brain tissue^31^. Based on this reasoning we hypothesize that the induction of tumor phenotype may have a biophysical basis. To test this hypothesis, we employed a well-established stochastic nanoroughness platform^25–27^ and investigated the effect of nano-scale physical cues on human fetal astrocytes. Our results demonstrate that physical cues can trigger phenotypic changes in astrocytes leading to the spontaneous formation of spheroids, which possess many of the hallmarks of GBM. Furthermore, these spheroids actively stimulate astrocyte migration and induce trafficking of human tumor cells.

## Results

### Astrocyte spheroid formation is triggered by nanoroughness and is independent of Piezo-1

The fate of primary human cortical astrocytes on substrates presenting stochastic nanoroughness (Rq) ranging from 12 nm and 32 nm (**Figure 1a**) over a 5-day period was investigated. The entire workflow for generating the substrates and the cell seeding is schematically described in **Supplementary Figure 1.** While astrocytes on the smooth substrate (Rq_3.5_) showed the classical *in vitro* astrocyte morphology, on substrates with a Rq of 12 nm, a spontaneous condensation of astrocytes was observed, with progressively increasing Rq’s yielding less condensed structures (**Figure 1b**). However, on day 3 and beyond, while the astrocytes on Rq’s of 16, 24, and 32 nm progressively dissociated to single cells, the spheroid on Rq of 12 retained their 3D-organization and underwent further maturation (**Figure 1b**). After 5 days of culture, the spheroids formed on Rq_12_ were further characterized using scanning electron microscopy (SEM) providing evidence for a three-dimensional structure composed of aggregated cells (**Figure 1c**). Since the substrate surface was stable over the duration of the study as verified using SEM (**Supplementary Figure 2**), and since physically masking surface of the substrates with Geltrex (commercial basement membrane matrix mix) coating abrogated spheroid formation on Rq_12_ surface and yielded monolayers of astrocytes by day 5 regardless of the substrate, the source of the physical trigger for spheroid formation can be attributed to the stochastic nanoroughness. In order to probe the role of cell proliferation in the observed phenomenon, cell numbers were quantified at day 5. Although astrocytes proliferated on all surfaces to different degrees, Rq of 12 showed the least proliferation with increasing Rq correlating with increased proliferation revealing a direct connection between nanoroughness and proliferative status of astrocytes (**Figure 1d**). Furthermore, since similar numbers of cells were seeded on each substrate at the beginning of the experiment, these findings indicate that low degrees of nanoroughness not only induce spheroid formation, but also reduce the proliferative capacity of astrocytes, as the initial attachment was not modulated by roughness (**Supplementary Figure 3)**. Since it is well established that proliferation of cells in a spheroid is usually lower than in monolayer, a vital dye was used which confirmed that the majority cells were viable refuting the existence of a necrotic core at the center of the spheroids (**Figure 1e**). Taken together, these findings demonstrate that astrocytes precisely sense and respond to stochastic nanoroughness and this directly influences the behavior and morphology of astrocytes. Besides the observed morphological changes, expression of EAAT2 which is responsible for 90% of the glutamate uptake in the adult CNS^32^ and GFAP, an intermediate filament protein considered a marker of astrocyte reactivity were investigated (**Figure 1f**). Both EAAT2 and GFAP expression showed no differences in comparison to controls implying that the astrocytes exposed to nanoroughness retain prominent physiological traits. In order to gain some mechanistic insights into how astrocytes perceive the underlying nano-scale physical cues, the overall expression patterns of the stretch-activated ion channel Piezo-1, which was previously demonstrated to be involved in sensing of stochastic nanoroughness by neurons^27^, and integrin αVβ3, because of its wide expression and implication in many physiological processes and diseases, were probed. Neither inhibition of Piezo-1 with GsMTx4, a potent inhibitor of piezo1, known to reversibly inhibit more than 80% of the mechanically induced currents^33^, nor blocking antibody towards αVβ3^34, 35^ had any effect on spheroid formation. (**Supplementary Figure 4**) suggesting that astrocytes perceive nano-scale cues through mechanism different from neurons.

**Figure 1:**
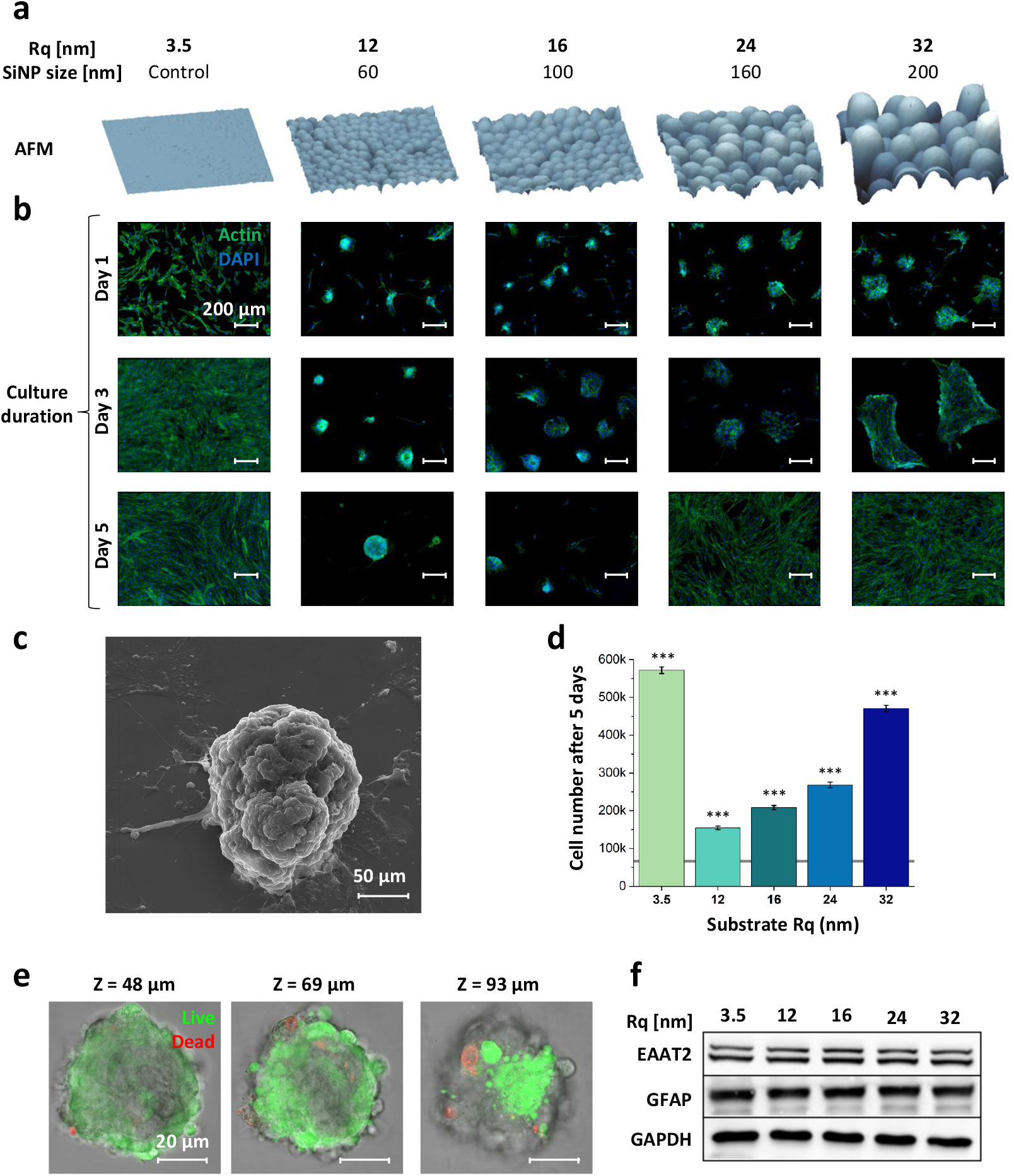
Nanoroughness alters astrocyte phenotype and induces spheroid formation. **(a)** AFM images of surfaces. **(b)** Astrocytes grown on varying degrees of nanoroughness show altered morphology revealed using actin staining. **(c)** SEM picture of astrocytes grown on Rq_12_ for 5 days, which aggregated into 3D spheroids. **(d)** Number of cells found on each substrate after 5 days of culture, obtained by DNA quantification (n = 3, ***p < 0.001). **(e)** Confocal imaging of a living astrocyte spheroid after 5 days of culture on Rq_12_, at three different heights. Live and dead cells are stained in green and red, respectively. **(f)** Western blot profile of GFAP and EAAT2 as function of exposure to nanorough. GAPDH served as the internal housekeeping control.

### Astrocyte spheroids remain stable even upon removal of physical cues and serve as aggregation centers for naïve astrocytes

Since scar formation by even highly reactive astrocytes generally is assumed to be permanent, and has been shown to be reversible under certain physiological conditions^36^, the stability of the astrocyte spheroids induced by nanoroughness was examined. To this aim, astrocyte spheroids matured for 5 days on Rq_12_ were trypsinized without impacting their integrity and transplanted on to glass substrates which represent an almost smooth surface (Rq 3.5 nm) (**Figure 2a**). Despite being the trigger for the astrocyte spheroid formation, withdrawal of the Rq_12_ nanoroughness did not lead to spheroid disintegration. Baring a slight reduction from the initial size, that can be attributed to the transplantation process, astonishingly, the spheroids remained intact and stable in size even after 30 days, implying that the phenotype induced by nanoroughness represents a permanent and fundamental transformation (**Figure 2b**). The fact that increasing astrocyte seeding density resulted in proportional increase in spheroid size with a dramatic (20-fold) increase above 270k cells/cm^2^ (**Figure 2 c and d)** led us to hypothesize that the spheroids are not derived from a single cell, but are the result of aggregation of distinct astrocytes which have undergone active association driven by a few astrocytes acting as aggregation centers. To test this hypothesis, an experiment was designed as schematically shown in **Figure 3a**: astrocytes were first transfected to constitutively express the fluorescent proteins BFP and TdTomato, allowing for simultaneous and independent tracking of both populations. BFP-expressing astrocytes were seeded on a Rq_12_ substrate for 5 days to induce spheroid formation and then, astrocytes expressing TdTomato were added to the substrates, and the evolution of the spheroids was monitored over a 5-day period using fluorescence microscopy. BFP-astrocytes formed spheroids as anticipated verifying that the transfection did not influence the spheroid formation. Following addition of TdTomato astrocytes, after 5 days, while Td-astrocytes also formed spheroids by themselves, many of the Td-astrocytes of fused with the *a priori* formed BFP-astrocyte spheroids, with 100% of BFP-spheroid showing the incorporation of TdTomato astrocytes and not vice versa (**Figure 3b**). In one such fusion event captured in **Figure 3c**, an aggregate of TdTomato expressing astrocytes can been seen collectively migrating a distance of 50 µm in a matter of 8 hours in order to fuse with the existing BFP expressing spheroid (**Supplementary Video 1**), suggesting that the colocalization resulted from direct attractive paracrine signals and random association, and that the established spheroid is stably anchored and serves as foci for the aggregation.

**Figure 2:**
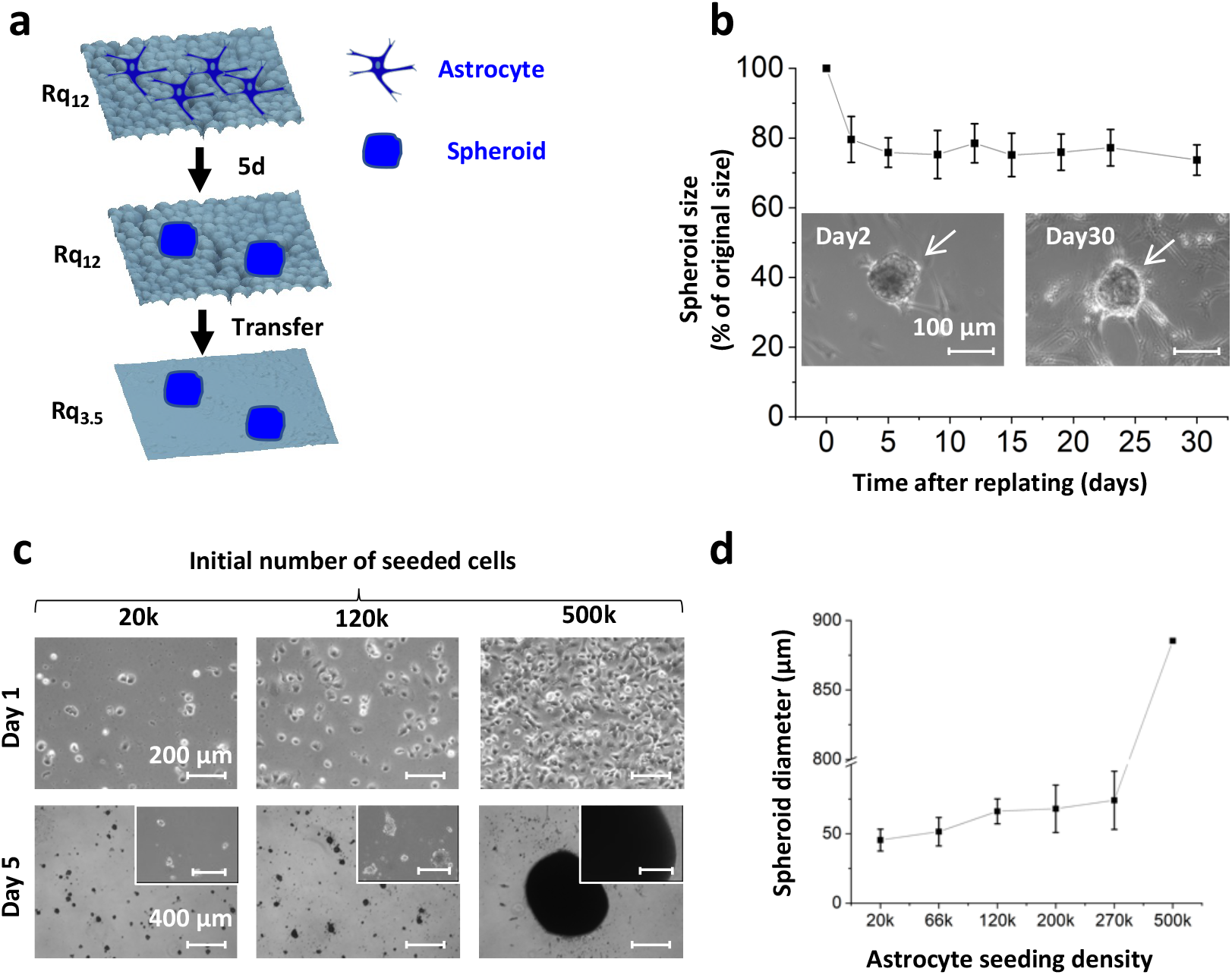
Stochastic nanoroughness-induced astrocyte spheroids have stable phenotype. **(a)** Schematic representation of the replating of astrocytes formed on Rq_12_ onto a smooth Rq_3.5_ substrate in panel b. **(b)** The size of spheroids formed on Rq_12_ replated on Rq_3.5_ remained the same for at least 30 days (n = 9). **(c)** Increase of the initial number of seeded cells resulted in the formation of larger spheroids **(d)** Quantification of the spheroid diameter with increasing seeding density, showing that initial seeding number is positively correlated with the final spheroid size. Error bars represent standard deviation from the mean (n=9).

**Figure 3:**
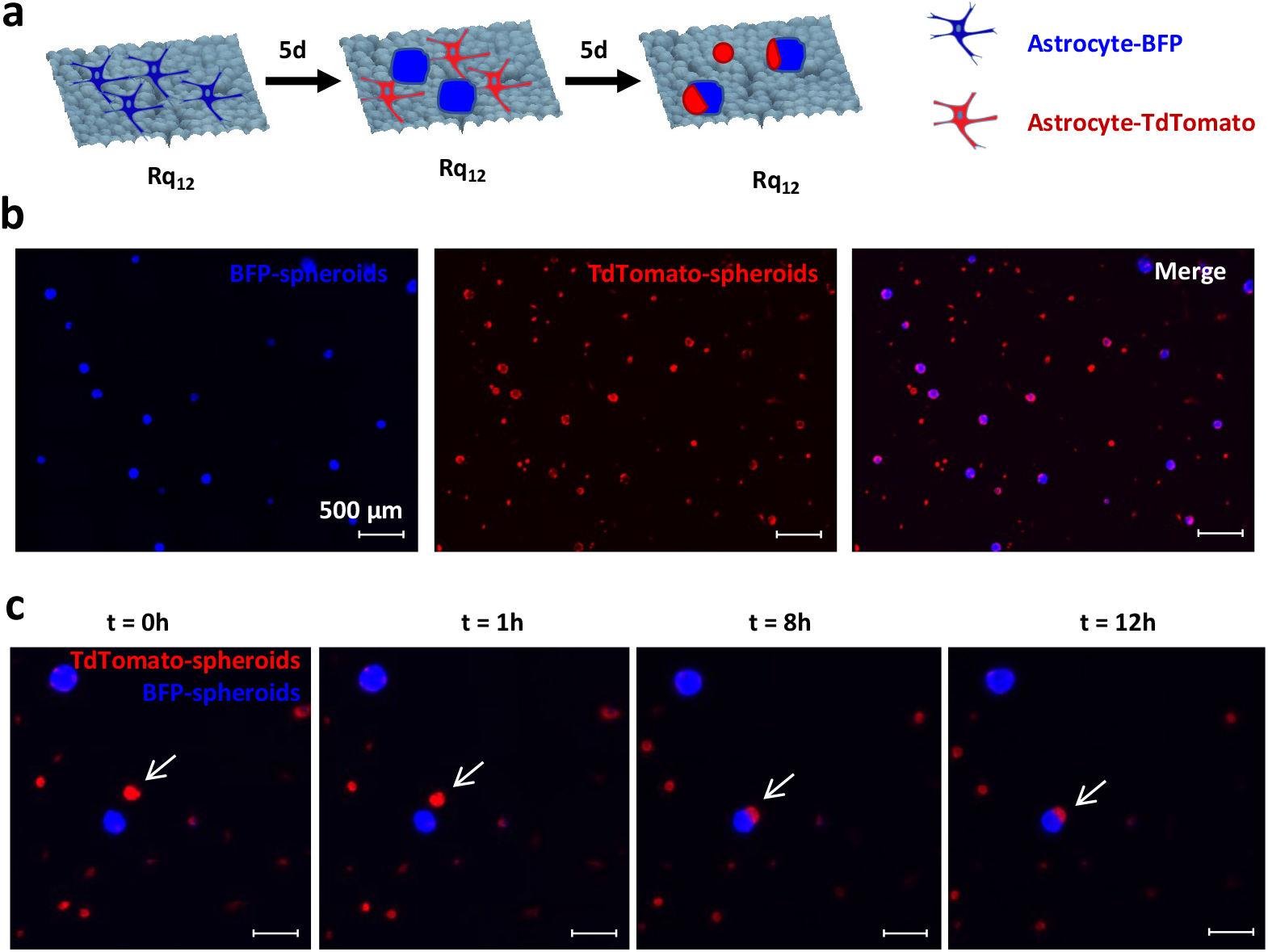
Spheroids function as aggregation centers for naïve astrocytes. **(a)** Schematic of the experiment design. **(b)** Five-days post-addition of TdTomato expressing astrocytes, 100 % of BFP fluorescence colocalize with TdTomato fluorescence signal demonstrating the incorporation of TdTomato astrocytes into the pre-formed BFP-spheroids. **(c)** Time course captured on day 4^th^ over 12 hours of a fusion event showing the collective migration of an aggregate of TdTomato astrocytes towards a pre-existing BFP spheroid leading to a fusion and incorporation of the TdTomato astrocyte population into the BFP spheroid.

### The gene expression patterns of astrocytes grown on Rq_12_ emulate those of tumoral cells

Since pathological conditions are often caused by dysfunction of multiple genes and gene networks, a global gene expression study using bulk RNA-seq was performed using the workflow shown in **Supplementary Figure 5**, allowing us to gain whole genome differential gene expression without bias. RNA extracted from astrocytes cultured on Rq_3.5_, Rq_12_, Rq_16_, Rq_24_ and Rq_32_ for 5 days i.e., the time when marked phenotypical changes are observed and well established, were analyzed. 2318 differentially expressed genes (DEGs) (p < 0.05) were identified in cells that were exposed to nanoroughness when compared to the cells grown on smooth substrate (Rq_3.5_), as visualized in a heatmap and Venn diagram (**Figure 4a, 4b**). Among these DEGs, 514 genes (22.17%) were fully specific to Rq_12_ (**Supplementary Figure 6**), which is the highest proportion among the four nanorough conditions. The DEG on Rq_16,_ Rq_24_ and Rq_32_ and other compartaive intersections are shown in **Supplementary Figure 7-12**. Unbiased hierarchical clustering grouped the samples into two groups with similar features. While, two extremes of the roughness studies - Rq_3.5_ and Rq_32_ - were close to each other, they differed significantly from Rq_12_, Rq_16_ and Rq_24_. This was confirmed by the principal component analysis (PCA) showing that the effect of stochastic nanoroughness, accounted for 75% of the observed differences, while experimental covariates and batch effects accounted for only 10% of the differences, suggesting the absence of “hidden effect” on the data set (**Supplementary Figure 13**). Taken together, this data matches the earlier phenotypical observations, and emphasizes the specificity of the phenotype of astrocytes forming spheroids on Rq_12_. Interestingly, among the 514 DEGs on Rq_12_, a majority of them (62%) was down-regulated. Based on the results of a study showing that a majority of down-regulated genes is a marker of dedifferentiation characteristics of a malignant transformation in a cellular model^37^, we propose that the astrocytes cultured on Rq_12_ might have undergone a transformation towards a cancerous phenotype. Further analysis of the 514 DEGs individually using UniProt^38^, enabled the manual sorting into functional categories (**Figure 4c**). Although it is challenging to determine the influence of individual genes on a biological mechanism, especially when genes within the same category show both down- and up-regulation, general trends could be deduced. Extracellular matrix (ECM) molecules were generally down regulated. Moreover, the balance of the expression of genes responsible for vesicular trafficking appeared to be directed towards an increase of the extracellular vesicle secretion, known to be one of the mechanisms by which cancer cells communicate with their surroundings^39^. Several genes involved in alternative splicing were downregulated, while none were up-regulated. This is a significant finding as the brain tissue is known to exhibit one of the highest degree of alternatively spliced transcripts, expressing a unique set of isoforms^40^, and reprograming of alternative splicing is a hallmark of cancer, where the tumoral cells express new specific splicing isoforms^41–43^. Also, a total of 14 genes involved in DNA repair were found to be downregulated, likely causing genome instability which is a characteristic of almost all human cancers^44^. Tumor suppressors overall were downregulated, including novel tumor suppressor candidates found in cervical and gastric cancer; VILL^45^, CDK5RAP3^46^, and KIAA0141^47^, along with the decrease in the expression of tumor protein P53 (TP53). TP53 mutations, leading to p53 loss were found to be the most frequent and earliest detectable genetic alteration in low-grade astrocytomas and GBM^48^ and contributes to glioblastoma progression^49^. Furthermore, TP53 is necessary to maintain a baseline expression of a wide range of tumor suppressor genes^50^. Not only are alterations of p53 visible in the cancer cells directly, but are also present in astrocytes at the tumor borders, favoring cancer malignancy^51^. Furthermore, several other genes implicated in cancer pathology, namely, TP53 inducible protein-11 (TP53I11)^52^, MTA1^53^, GLIPR1, GNA13^54^, NCEH1^55^, GMCL1^56^, and BEX2^57^, were differentially expressed.

**Figure 4:**
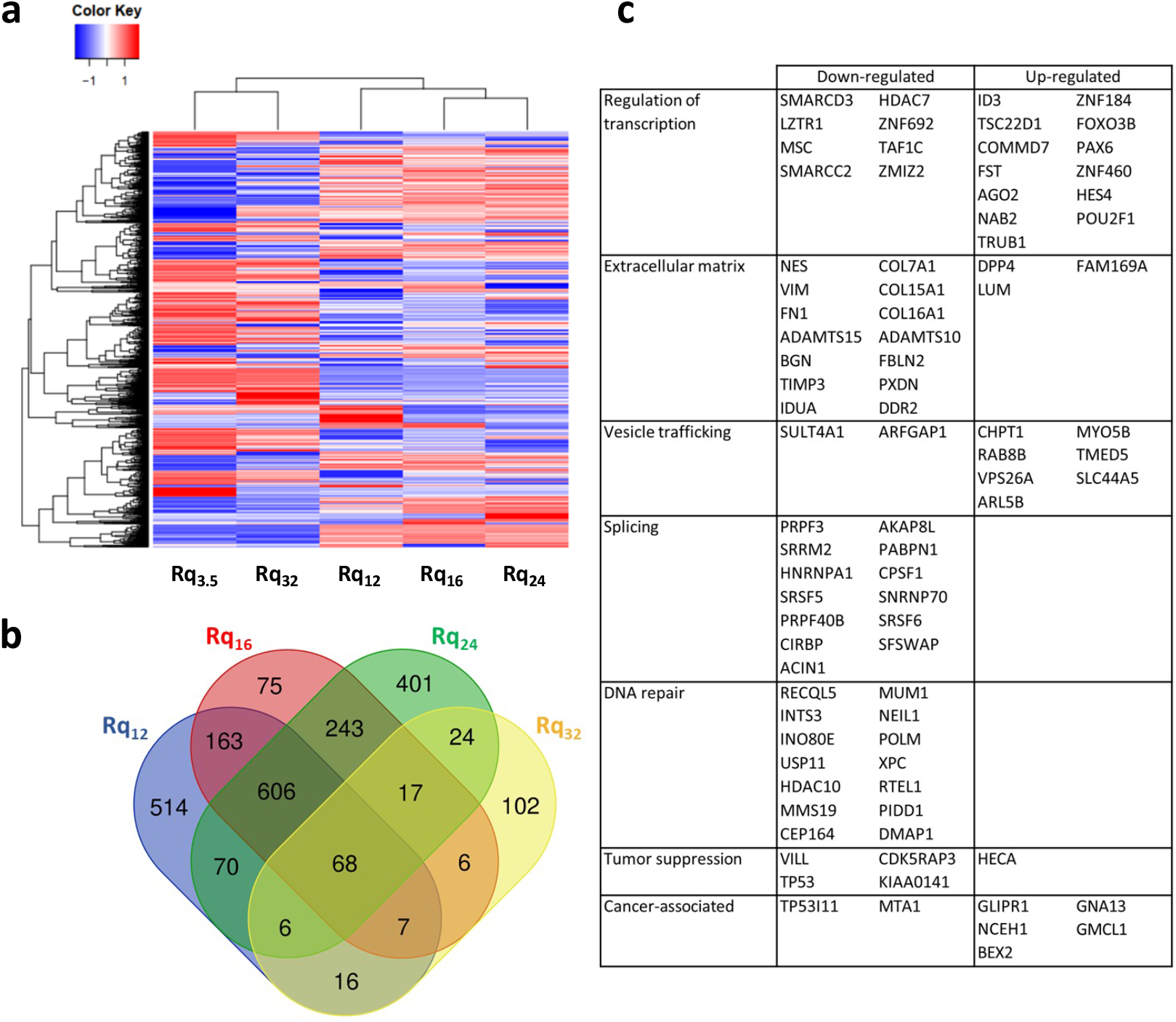
Global gene expression of astrocytes is altered by stochastic nanoroughness. **(a)** Heatmap representing the 2318 DEGs in the different culture conditions compared to astrocytes cultured on Rq_3.5_, which is glass substrate served as the reference. The black lines on top of the heatmap represent the clustering of the different conditions, highlighting the similarity between the extremes of the roughness range, and pointing the uniqueness of the Rq_12_ spheroidal phenotype in terms of gene expression (n=2, the average result is shown). **(b)** Venn diagram showing the overlap between the DEGs and the culture conditions. Astrocytes grown on Rq_12_ are unique with 514 DEGs being specific to this condition. **(c)** Classification of the 514 DEGs specific to Rq_12_ into functional categories related to the cancer pathology.

In order to fully appreciate how the selected genes, work in concert to drive cancer-related processes, the genetic interactions between the DEGs were studied, allowing for a statistical assessment of the relationships between genes. The meta-database String (version 11.5) was used to map all known evidences of interactions between genes to establish a gene interaction network (**Figure 5**)^58^. According to the String analysis, with an average node degree of 1.18 and an enrichment p-value of 0.00813, the DEGs from Rq_12_ were found to have more interaction among themselves than what would be expected from a random set of proteins of similar size drawn from the genome. With an increase of the nodes by 20%, such an enrichment indicates that the proteins are at least partially biologically connected as a group. A big node was evidenced, centered around the TP53. It is noteworthy that it has been shown that cancer cells transform neighboring astrocytes by reducing their p53 expression, thus shutting down the cell-death pathways in both cell types, favoring the tumor progression^51^. String analysis further revealed three prominent signaling nodes involving ADAMTS proteases, NOTCH3 and fibronectins, suggesting that these set of genes maybe integral to the phenotype observed in the spheroids.

**Figure 5:**
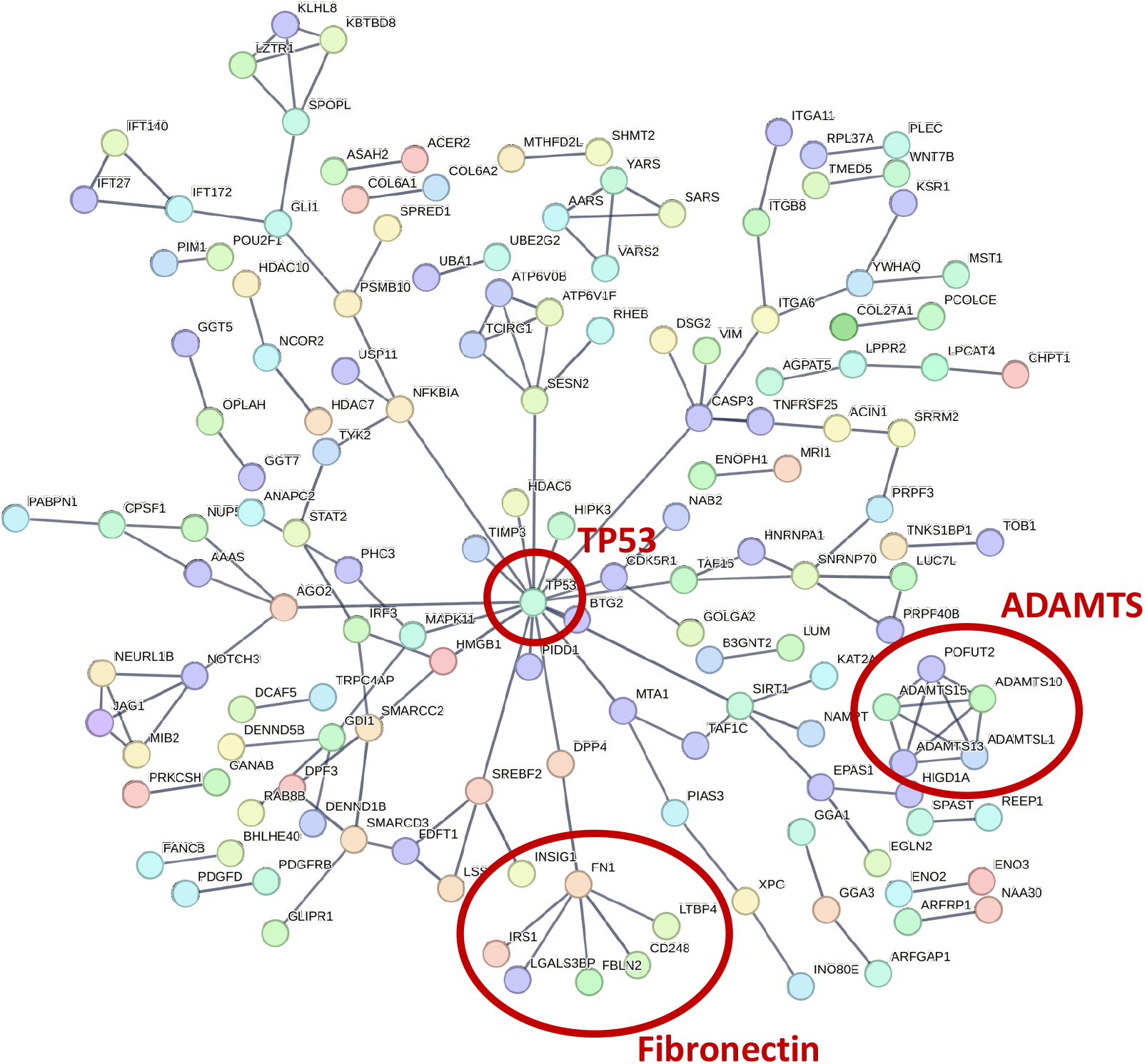
Gene interaction network analysis of DEGs by nanoroughness-induced astrocyte spheroids: String analysis of the 514 DEGs specific to Rq_12_ revealing gene expression patterns similar to glioma cells involving TP53 locus, with further involvement of NOTCH3 and fibronectin loci.

### Stochastic nanoroughness confers cancer stem cell-like characteristics to astrocytes

Since senescence, the arrest of the cell-cycle, crucial during embryonic development and tissue remodeling including wound-healing has been implicated in cancer and plays a role in preventing the proliferation of cells experiencing oncogenic stress^59^, we investigated whether nanotopography could trigger senescence. Astrocytes were cultured on the different roughness regimes, and were stained for β-galactosidase (β-gal), a known marker for senescence^60^ and are also expressed by aging hippocampal neurons^61^. Spheroids formed on Rq_12_ were strongly positive for β-gal suggesting that they have undergone senescence, which was further confirmed by the lower percentage of Ki67-positive cells (**Figure 6a and 6b**). The induction of senescence is consistent with the observation that spheroid size remained constant after replating **Figure 2b**. Since senescent cells are known to display a unique senescence-associated secretory phenotype (SASP) which has potentially detrimental long-term implications^62^, the release of the cytokine IL6 was investigated. While higher IL-6 levels were observed in all nanoroughness conditions a significantly higher IL6 production per cell was found in the Rq_12_ condition, characteristic of SASP (**Figure 6c**). This is a significant finding as this is the first evidence of astrocyte senescence triggered by physical cues as opposed to environmental triggers such as oxidative stress, radiation, hyperoxia, oncogenes (RAS or Raf) and replicative exhaustion^63–66^.

**Figure 6:**
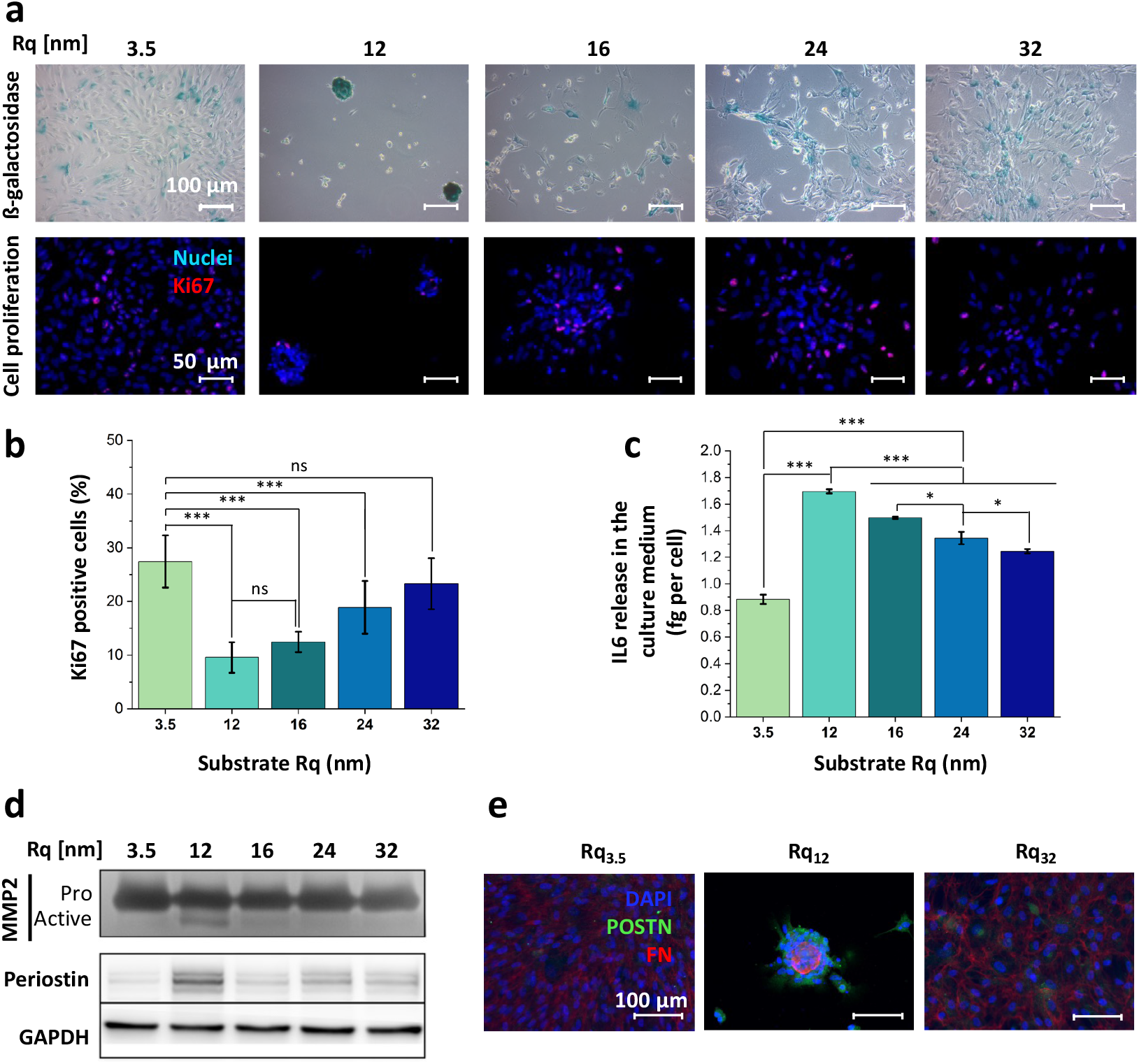
Nanoroughness induces cancer cell characteristics in astrocytes. Staining for β-galactosidase (β-gal) and Ki67, two markers of cellular senescence revealing a higher β-gal postivity **(a, top panel)** and dimninished proliferation capacity in strocytes grown on Rq_12_ **(a, bottom panel and b)** (Rq_3.5_, n = 8, Rq_12_, n = 11, Rq_16_, n = 4, Rq_24_, n = 9, Rq_32_, n = 7). **(c)** Astrocytes grown on Rq_12_ release more IL6, characteristic of the senescence-associated secretory phenotype (n = 3). **(d)** Astrocytes grown on Rq_12_ show activation of MMP2 revealed by zymography, along with an increase in periostin (POSTN) expression, shown by western blot. GAPDH was used as an internal loading control (pooled extracts from n=3). **(e**) Astrocytes grown on Rq_12_ express more periostin than their counterparts grown on Rq_3.5_ and Rq_32_, as shown using immunofluorescent staining. (*p < 0.05, **p < 0.01, ***p < 0.001, ns no statistical difference).

Matrix metalloproteinases (MMPs) expression is dysregulated in several pathological conditions, particularly cancer where they have been one of the most researched therapeutic targets and biomarkers^67^. Since senescent cancer-associated fibroblasts have been shown to secrete pro-MMP-2^68^, and in the CNS, MMPs are mainly produced by astrocytes particularly in response to injury-related activation^69^, the influence of nanoroughness on astrocytic release of MMP-2 was investigated. Although roughness regimen did not influence pro-MMP2 release, active-MMP-2 was uniquely detected in the culture medium of astrocytes grown on Rq_12_ (**Figure 6d**). A positive correlation between astrocyte seeding density and the amount of activated MMP-2 evidenced that astrocytes are directly involved in the activation process (**Supplementary Figure 14**) and demonstrating MMP-2 activation as consequence of astrocyte senescence. To date, this is the first evidence of MMP-2 activation by astrocytes directly triggered by physical cues which might involve astrocyte senescence, and provides further evidence for brain ECM topography as one of the contributing factors in neurodegeneration^27^. Since MMP-2 can localize on the surface of invasive cells via integrin binding and promote cell-mediated collagen degradation^70^, reactive astrocytes have been detected around gliomas^13, 71^, and it has been shown that activation of MMP-2 secreted by astrocytes require interaction with glioma cells^72^, we suggest that the ability of senescent cells to shape the microenvironment is not limited to cytokine and protease secretion, but may also include differential ECM production. We therefore investigated the secretion of periostin (POSTN) a matricellular ECM protein with known role in tissue remodeling^73^, and cancer progression and metastasis^74, 75^ whose expression in cancers is associated with poor prognosis. Western blot analysis showed a strong upregulation of POSTN by astrocytes cultured on Rq_12_ (**Figure 6d**) and this was also confirmed by immunohistochemistry (**Figure 6e**). Spheroids induced by Rq_12_ showed prominent staining for POSTN which was uniform throughout the entire spheroid (**Supplementary Figure 15**), while in comparison astrocytes cultured on Rq_3.5_ astrocytes and Rq_32_ showed no such expression providing further evidence that astrocytes in the spheroid represent a unique phenotype. Since high POSTN expression has been observed in tissues (periosteum, periodontal ligament) subjected to constant mechanical stress, it is thought to play a role in mechanoreception, and its upregulation in astrocytes exposed to nanoroughness reinforce this postulated role^76, 77^. Furthermore, as POSTN expression is low under healthy conditions but upregulated following injuries^78^, the prominent expression of POSTN by astrocytes grown on Rq_12_ represents an hitherto unexplored marker for activated astrocytes.

### Stochastic nanoroughness, through induction of cellular senescence, mediates the crosstalk between astrocytes and cancer cells

Senescent tumor cells have been shown to lead collective invasion in thyroid cancer^79^, and MMPs secreted by senescent fibroblasts were found to support tumor xenograft growth^80^. Furthermore, senescent cells act as aggregation center for cancer cells, therefore participating in three-dimensional cluster formation^81^ through a mechanism involving the SASP. Building on these findings and our observation that spheroids serve as aggregation center for naïve astrocytes, we postulated that astrocyte senescence, mediated by nanoroughness, may provide attractive cues to invading or metastatic tumor cells. To test this, co-cultures of astrocytes with the GBM cell line U87 or the triple negative breast cancer cell line MDA-MB-231 were studied on surfaces of varying nanoroughness (**Figure 7a**). After 4 days of co-culture, U87 cells were homogenously distributed on Rq’s except on Rq_12_ where they formed aggregates around the astrocyte spheroids (**Figure 7b**). Similarly, after 16 days of co-culture, MDA-MB-231 cells also exhibited association with the astrocyte spheroids on Rq_12_ (**Figure 7c**) suggesting that in both systems the cancer cells are drawn toward the spheroid microenvironment. Since both U87 and MDA-MB-231 cells when cultured on their own over the duration of the experiment do not exhibit discernable changes in phenotypical traits due to nanoroughness (**Supplementary Figure 16**), the associative tendencies between cancer cells and astrocyte spheroids on Rq_12_ can be attributed to the astrocyte phenotype. Because breast cancer together with melanoma and lung cancer are the three tumor types known to metastasize to the brain^82^, we conclude here that astrocytes and nanoroughness may synergistically function as a specialized niche microenvironment to attract circulating tumor cells and promote metastatic lesions^83^. Our results also reinforce findings from a previous study showing the ability of astrocytes to alter migration and morphology of metastatic breast cancer cells^84^. In order to rule out that cancer cells are guided towards the astrocyte spheroids by ECM tracks, the opposite experiment was performed, where astrocyte spheroids formed on Rq_12_ were trypsinized and transferred onto U87 cells cultured on smooth Rq_3.5._ Here too, U87 cells migrated toward the astrocyte spheroids after 48-hours of co-culture, confirming the attraction of the cancer cell towards the senescent astrocyte population in the spheroids (**Supplementary Figure 17a and 17b**). This experiment being performed on a smooth (Rq_3.5_) substrate confirmed that the effect of nanoroughness on the crosstalk is rather indirect, by durable modification of astrocyte phenotype and SASP. The attraction of U87 cells to the astrocyte spheroids was however not accompanied by an increase in proliferation as assessed by MTS assay (**Supplementary Figure 17c**), suggesting that interactions between U87 and astrocytes are of an attractive nature and do not involve increased local proliferation, further proving that the SASP astrocyte phenotype exhibits unique crosstalk with cancer cells. These findings are highly significant as it for the first time presents evidence for a biophysical basis for induction of oncogenic phenotype in astrocytes and provides the impetus to further investigate the role of local changes in ECM accumulation and aggregation in induction of aberrant phenotype in astroglial population. In a preliminary experiment, we have shown that astrocytes dissociated from spheroids grown on Rq_12_ after stereotaxic implantation in mice brains remarkably survive for over 12 months and disseminate (migrate) away from the site of injection (**Supplementary Figure 18**). The human origin of these cells was confirmed by Alu-repeat staining which is specific for human nuclei. Future experiments will aim at co-injecting astrocytes grown on the different nanorough substrates with GBM cells to assess whether the astrocytes grown on Rq_12_ possess the ability to facilitate tumor progression *in vivo*.

**Figure 7:**
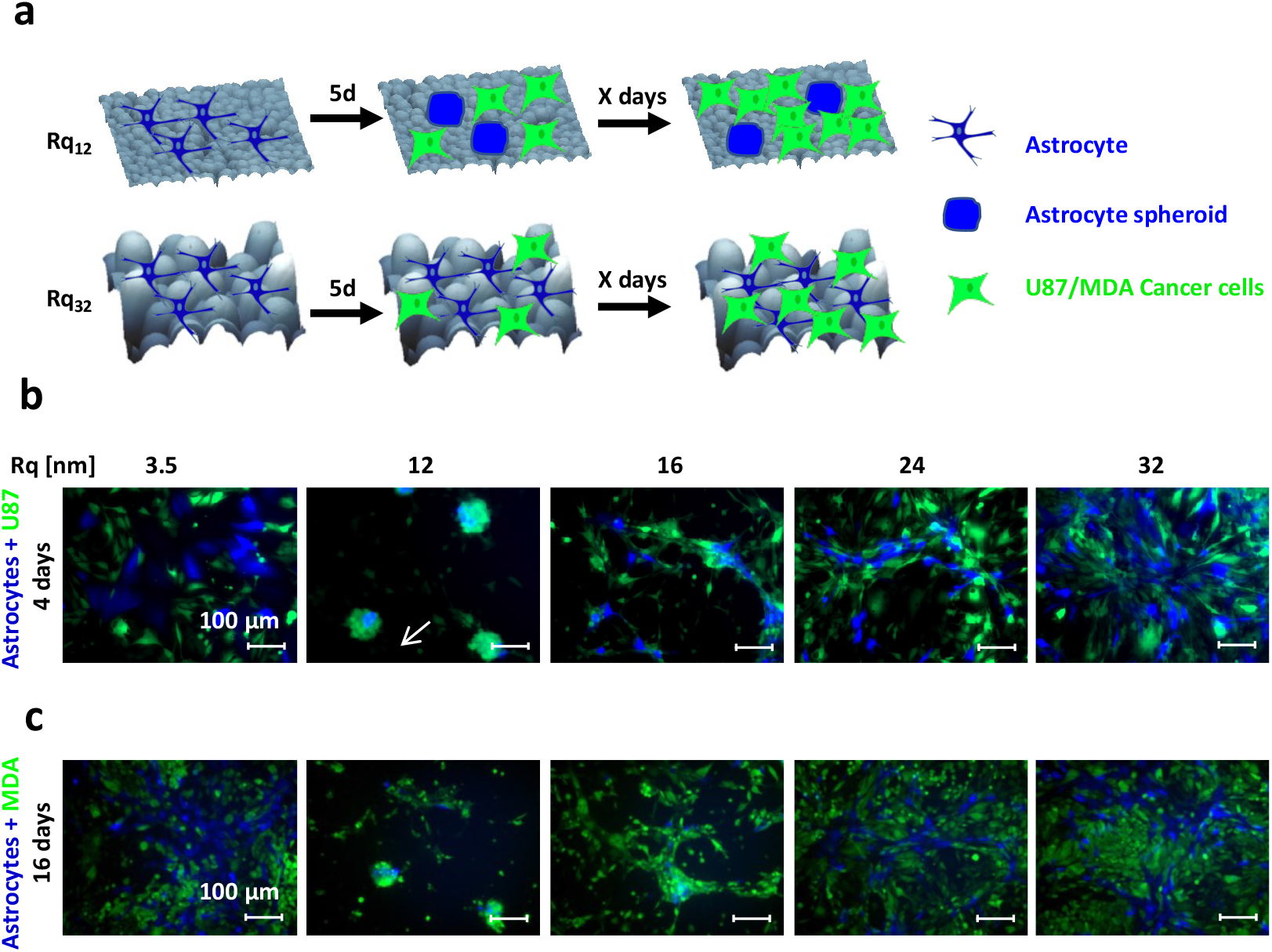
Stochastic nanoroughness mediates the interactions between astrocyte and cancer cells. **(a)** Schematical representation of the experimental design. Naïve astrocytes were cultured for 5 days to establish the respective phenotype on varying stochastic nanoroughness and then cancer cells U87 or MDA-MB-231 were seeded and their fate was followed until clear trends emerged. **(b)** Fluorescence images after 4 days of co-culture of BFP expressing astrocytes and green fluorescent protein (GFP) expressing U87 cells and, **(c)** Fluorescence images after 16 days of co-culture of BFP expressing astrocytes and GFP expressing MDA-MBA-231 cells.

## Discussion

Our findings that stochastic nanoroughness can alter naïve astrocyte phenotype and induce their spontaneous association into spheroids which possess cancer-like characteristics is quite significant as it provides evidence for a potential biophysical basis for the formation of glioma. Furthermore, the ability of spheroid derived-astrocytes to survive and migrate from the site of implantation in mice brain (**Supplementary Figure 18**) implies that astrocytes exposed to physical cues can persist in vivo and engage distant locations. When placed in context of our earlier findings that topography of the brain tissue is altered around Alzheimer plaques^27^, and that PGs contribute prominently to the mechanical properties of the brain^31^, the evidence presented here provides a compelling argument to consider changes to the microstructure of the brain in the induction of neurodegenerative conditions.

Among the many pro-oncogenic factors NOTCH (1-4) signaling is known to play an important role in tumorigenesis^85, 86^ and also in the progression and invasion of GBM^87^. NOTCH signaling is a highly conserved juxtracrine signaling paradigm that requires cell-cell contact^88^. NOTCH3 has been shown to correlate with poor prognosis and increased invasive traits in GBM^89^. We have shown that in breast cancer spheroids, the activation of mesenchymal-NOTCH signaling requires mesenchymal-endothelial crosstalk and this is mediated by the breast cancer cell phenotype^90^. The down regulation of NOTCH3 in the astrocyte spheroids suggests that the gaining invasive traits might require the acquisition of other cell types such as vasculature and immune cells to fulfill the juxtacrine signaling. In addition to soluble cues, extracellular matrix FN expression has been associated with progression^91^ and invasion of gliomas^92^, and gene ontology enrichment analysis has revealed that FN and ECM related genes are in general upregulated in GBM tissue versus healthy brain^93^. Here a downregulation in FN expression was observed in astrocytes within spheroids and this might suggest that other extraneous factors only accessible in the in vivo environment are necessary to drive an aggressive phenotype. Taken in sum, the gene interaction analysis reveals an astrocyte phenotype within the spheroids that bears many cancer-related traits. In future studies, we aim to elucidate the role of microglia, and crosstalk between endothelial cells derived from glioma environments and activated monocytes in consolidating the GBM-like phenotype and driving invasive traits in the astrocyte spheroids.

Considering the findings of this study, the idea of conditioned astrocytes (defined reactive astrocyte population) as a therapeutic tool in treating neurodegenerative conditions is worth considering. Towards this objective, a detailed analysis of the epigenome of astrocytes exposed or conditioned to biophysical cues could identify new targets for mitigating malevolent changes to astrocyte physiology. Furthermore, one could also envision implantation of astrocytes conditioned ex vivo using physical cues to bestow protective traits as means to predictably alter cellular and signaling homeostasis in the brain.

In conclusion, by elucidating the role of physical cues, our study provides novel insights into how the microstructure of the brain may play a pivotal role in modulating astrocyte behavior and contributes to a better understanding of the complex interplay between biophysical factors and soluble signals in tumor induction in the brain.

## Materials and Methods

### Cells

Human fetal astrocytes were purchased from Sciencell (#1800) and cultured in complete astrocyte medium from Sciencell including 2% FBS (#1801) in a humidified incubator at 37°C and 5% CO_2_. Cell adherence was ensured by coating with Poly-L-Lysine at 2 µg/cm^2^. Human glioblastoma-derived cell line, U87 was purchased from ATCC^©^ and cultured in DMEM + 10% FBS in a humidified incubator at 37°C and 5% CO_2_. Human triple negative breast cancer cell line MDA-MBA-231 cell line was purchased from ATCC^©^ and cultured in DMEM + 10% FBS in a humidified incubator at 37°C and 5% CO_2_. All cell lines were genotyped and tested negative for mycoplasma. Studies were replicated using two independent astrocyte donors. All experiments were performed using astrocytes at passage three.

### Transfection

Lentiviral particles containing BFP: pLVX-mTagBFP2-P2A-Puro (BIOSS Toolbox, University of Freiburg), GFP: pGIPZ (Openbiosystems, RHS4 346), and tdTomato: pLVX-tdTomato: (BIOSS Toolbox, University of Freiburg) were produced in HEK293 cells by mixing lentiviral vector and packaging vectors using branched polyethyleneimine (bPEI) (MW 25Kda, Sigma) as the transfection reagent. For transfection, 5 µg of DNA (4:3:1 of a transgene, pCMVdR8.74 (packaging plasmid; Addgene, Plasmid #22036) and pMD2.G (envelope plasmid, Addgene, Plasmid #12259) were diluted in 250 µL Opti-MEM (Invitrogen), afterwards, 11.25μL of bPEI (1 mg/mL) was added and incubated for 25 min at room temperature before transferring to HEK293 cells. 16 h after transfection, the medium was exchanged with the medium of the target cells. and 24 and 48 h after that, the media containing lentiviral particles were accumulated and filtered through a sterile 0.20 µm syringe filter (Millipore) to infect target cells. Infected cells include the human fetal astrocytes, U87 and MDA-MB-231 cell lines. Three days after transduction, infected cells were selected by adding puromycin (2 µg/mL) (Sigma) to the culture medium.

### Preparation of surfaces presenting stochastic roughness

Surfaces presenting stochastic nanoroughness were prepared by spin-coating silica (SiO2) nanoparticles (SiNP) on glass substrates (Supplementary Figure 1). Briefly, SiNP were synthesized using the Stöber process as described earlier^25, 27, 94^ via ammonia-catalyzed hydrolysis of tetraethylorthosilicate (TEOS) in 200 proof ethanol. SiNP size was tuned by varying the amount of ammonia and TEOS in the reaction, The size of the SiNP was confirmed using dynamic light scattering (DLS) analysis performed on DelsaNano C equipped with the DelsaNano Software V3.73/2.3 (Beckman Coulter Inc.) in triplicate. Then, glass slides of 5cm^2^ (2.5 x 2.0 cm) were cleaned by successive immersion in toluene – acetone – ethanol for 5 minutes each in a slide holder placed an ultrasonic bath, and following a drying period of 30min at 60°C, 200 µL of SiNP solution of the desired size was deposited at low-speed (5s, 400 RPM) followed by film casting (20s, 2000 RPM) using a spin coating (P6700 Series; Specialty Coating systems, Inc.; Indianapolis, USA). To ensure homogenous coatings seven layers were deposited and then the substrates were dried for 12 hours at 200°C. The presence of nanoroughness was confirmed using tapping mode on Dimension V Bruker AFM equipped with phosphorus silica doped cantilever (40N/m, 300kHz, Symmetric Tip), at a scan rate of 0.9 Hz with 256 scans per image. The root-mean square of the roughness (R_q_) was measured and is reported as an average of measurements at five different positions over an area of 1 µm^2^ at five different positions and calculated using NanoScope Analysis 1.20.

### Seeding of astrocytes on nanorough substrates

Substrates were placed in 6-well plates and sterilized for one hour under UV light, then coated with poly-ornithine (20 µg/mL; 4 µg/cm^2^) for 1h at 37°C followed by laminin coating (5 µg/mL; 1 µg/cm^2^) for 2h at 37°C. Astrocyte suspension (66000 cells/100 µL) was incubated on each substrate for 30 minutes to ensure optimal attachment, then 2 mL of astrocyte medium were added (**Supplementary Figure 1**).

### Microscopy

#### Immunofluorescence

Cells were fixed in 3.7% formaldehyde for 15min, and then blocked with 2.5% goat serum, 0.1% triton-X, 0.05% tween20 in PBS for 1h at room temperature. Three markers were stained using the following primary antibodies: Ki67 (Abcam, 1:500 dilution), POSTN (Abcam, 1:200 dilution) and Fibronectin (Santa Cruz Biotechnology, 1:200 dilution). Primary antibody was incubated overnight at 4°C, then secondary Alexa-conjugated antibody was incubated for 75min at room temperature. Cell nuclei were stained using 1:5000 Hoechst (Invitrogen) solution in PBS for 15 minutes at room temperature before samples were mounted in vectashield vibrance antifade (Vector labs).

#### Actin cytoskeleton

For visualization of the actin cytoskeleton, astrocytes were fixed in 3.7% formaldehyde for 15min, permeabilized in 0.1% Triton-X in PBS for 15min, and incubated in 165 nM solution of Phalloidin conjugated with fluorochrome Alexa-488 (Invitrogen) for 30min. Cell nuclei were stained using 1:5000 Hoechst (Invitrogen) solution in PBS for 15 minutes at room temperature before samples were mounted in vectashield vibrance antifade (Vector labs).

#### Scanning electron microscopy

Scanning electron microscopy images of cells on substrates were obtained as follows: Cells were fixed in 2.5% glutaraldehyde for 30min, washed in PBS, and dehydrated by incubation for 3min in each of the increasing ethanol concentrations (10%, 20%, 40%, 60%, 80%, 100%). After an overnight drying period, fixed cells on their glass substrate were coated with a gold layer prior to their imaging in a Quanta 250 FEG equipped with the FEI x T software.

#### Live-Dead staining of spheroids

Full astrocyte spheroids were stained for live and dead cells by incubation in 2 µM calcein AM and 5 µM ethidium homodimer-1 solution for 30min followed by immediate imaging using a confocal ZEISS LSM 880 Observer equipped with the software ZEN Black 2.3.

#### β-Galactosidase staining

Staining for b-galactosidase was carried out using a commercial kit (Cell Signaling Technology) following manufacturer’s instructions. Astrocytes were fixed for 15 minutes in the supplied fixing solution, then incubated with the staining solution adjusted to pH = 6 overnight at 37°C, before imaging.

##### ELISA for IL6

Conditioned media were collected after 5 days of culture and centrifugated at 600g for 10 minutes to remove cell debris. IL6 concentration was measured using ELISA commercial kit according to manufacturer instructions (R&D Systems, #DY206).

### RNA-sequencing

The source RNA for the sequencing runs was extracted using the RNeasy Micro kit (Qiagen) and nucleic acid concentration was measured using a Nanodrop 2000c. RNA was used to prepare poly-A selected directional libraries using the NEB Next Ultra Directional RNA Library Prep Kit (New England BioLabs, USA) and libraries were sequenced utilizing the Illumina Hiseq2000 platform at the sequencing core facility of the Max Planck Institute of Immunology and epigenetics (Freiburg, Germany). Paired-ends reads generated from sequencing on the Illumina Hiseq2000 were analyzed using the open web-based Galaxy platform^95^. An Overview of the RNA-Seq computational analysis pipeline can be found in Supplementary Figure 11.

### Western blot

Proteins were extracted from astrocytes lysed with radioimmunoprecipitation assay (RIPA) buffer *and were* quantified using the Pierce BCA assay (ThermoFisher). Proteins (11µg/well, boiled 95°C/5min) were separated through electrophoresis on an 8% polyacrylamide gel in denaturing conditions. Transfer of proteins onto PVDF membranes (Bio-rad) was performed for 75 minutes at 100V. Membranes were then blocked in 5% BSA in TBST for 1 hour at room temperature. Four markers were assayed using the following primary antibodies: POSTN (Abcam, 1:200 dilution), GFAP (Novus Biologicals, 1:1000 dilution), EAAT2 (Santa Cruz Biotechnology, 1:100 dilution) and GAPDH (Santa Cruz Biotechnology, 1:500 dilution). Primary antibodies were incubated overnight in 2.5% BSA in TBST, followed by incubation with HRP-conjugated secondary antibodies at 1:2500 in 2.5% BSA in TBST for 75 minutes at room temperature. Bands were revealed using SuperSignal West Pico PLUS chemiluminescent substrate, and imaged using Fusion FX7 (Vilber).

### Zymography

Conditioned media were collected and centrifugated at 600g for 10 minutes to remove cell debris, then subjected to non-denaturing electrophoresis in 8% polyacrylamide gel containing 1mg/mL gelatin. Gels were washed two times in 2.5% Triton-X for 30min, then incubated 24 hours in SimplyBlue SafeStain at room temperature, before being imaged with a Fusion FX7 (Vilber) imaging device.

### Cell DNA quantification

Cells were lysed in a 0.5% Triton X-100 and 20 mM ammonium hydroxide (NH_4_OH) buffer. DNA was quantified using the Quant-iT™ PicoGreen™ dsDNA Assay Kit (ThermoFisher) read using a Synergy HT plate reader (Biotek). The number of cells was calculated considering 6 pg DNA per cell.

### Orthotopic implantation and brain collection

Orthotopic implantation of the cells and spheroids was carried out as described previously^96^. Mice were anesthetized with 2.5% isoflurane in an induction chamber. Anesthesia was maintained at 1.5% isoflurane delivered through a nose adaptor on the Neurostar stereotactic frame (Neurostar, Tübingen, Germany). A midline incision was made and a burr hole was drilled at 1 mm posterior to the Bregma, and 2 mm lateral from the midline to the right. Cells were dissociated from the spheroids by trypsinization and stereotactic implantation of the cells (100,000 cells in 4µL PBS) was performed using a Hamilton 10 µL syringe (#80300, 701N, 26s/2”/2, Hamilton) at a depth of 3mm below the dura surface. Two minutes after injection ended, the needle was slowly retracted to avoid reflux of the cell suspension. The scalp wound was closed with sutures (5-0, Polypropylene suture, Ethicon, USA). Twelve months later, animals were transcardially perfused with ice cold PBS and brains were dissected, fixed in formalin and embedded in paraffin for in situ hybridization. Staining of human nuclei was carried out by chromogenic *in situ* hybridization (Zytovision kit) for human Alu repeat sequences following the manufacturer’s instructions. Nuclear fast red (Sigma) staining was used as nuclear counterstaining.

### Statistical analysis

All values are presented as the average ± standard deviation of the mean. Unpaired t-test provided in the Microsoft excel statistical analysis package was used to determine statistical significance. P-values ≤ 0.05 were considered statistically significant.

## Acknowledgments

The authors wish to thank Francine Wolf for the Alu-staining and Dr. Tala Shekarian for assistance with mice implantation.

## Author contributions

Conceptualization: VPS, LS

Methodology: LS, MS, RS

Investigation: LS, MS

Resources: VPS, GH

Visualization: LS, MS

Supervision: VPS, BH

Writing—original draft: LS, VPS

Writing—review & editing: LS, MS, RS, BH, MR, GH, VPS

## Data and materials availability

All data are available in the main text or the supplementary materials.”

