## Supplementary Information for "Biophysical basis for the induction of glioblastoma-like phenotype in astrocytes"

### **Supplementary Figures** (Figs. S1 to S18)

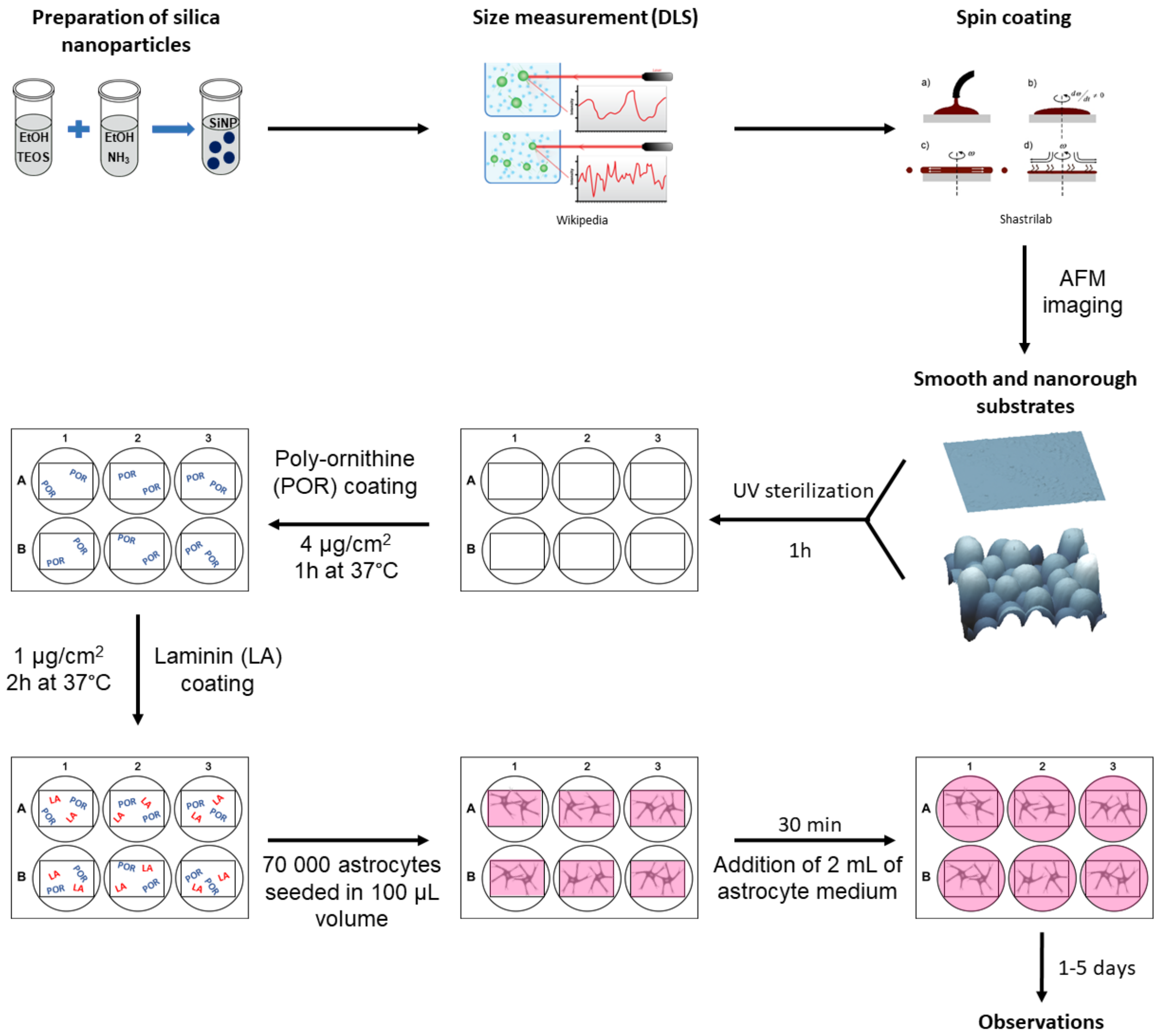


**Fig. S1.** Experimental design. Silica nanoparticles of various sizes were synthesized using the Stöber process, measured using dynamic light scattering, spin coated on glass substrates, and stochastic nanoroughness (Rq) was assessed using AFM. For cell culture, the prepared nanorough substrates were sterilized, coated with Poly-ornithine and laminin, and then seeded with astrocytes.

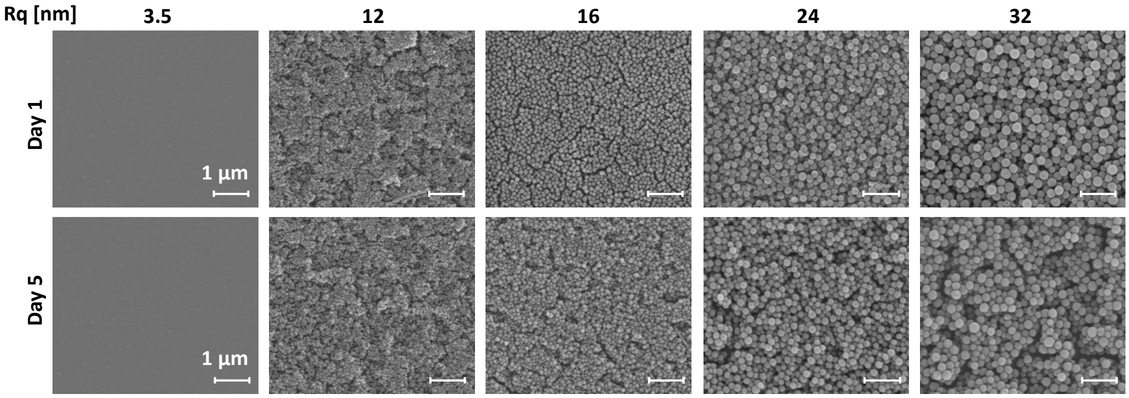


**Fig. S2.** Nanorough substrates remain stable under the cell culture conditions, at 37°C.

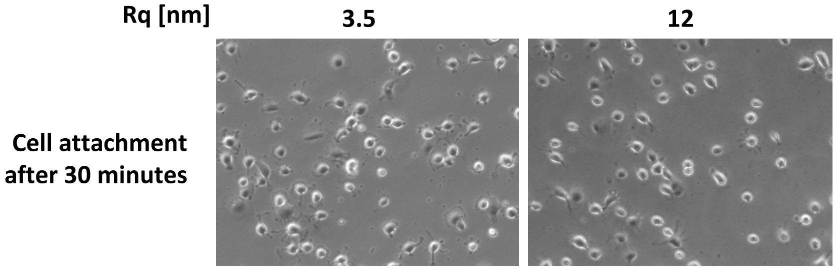


**Fig. S3.** Initial cell adhesion is not impacted by nanoroughness. Astrocytes were imaged 30 minutes after seeding. 68 cells were counted on $Rq_{3.5}$, while 67 were counted on $Rq_{12}$.

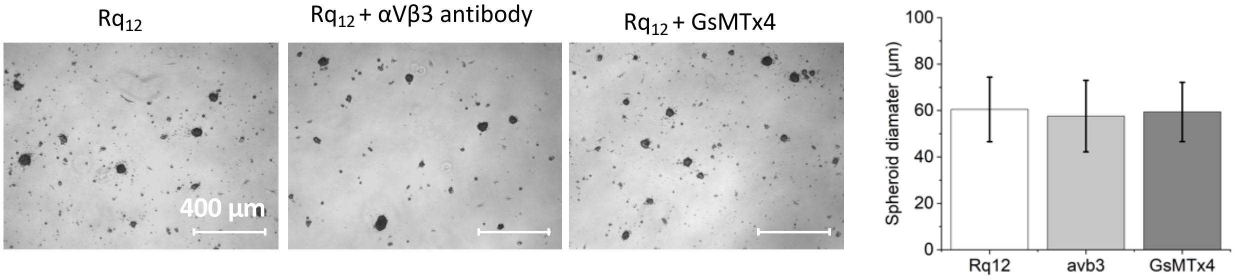


**Fig. S4.** Blocking integrin αVβ3 using a blocking antibody, or blocking the piezo1 receptor using the spider venom toxin GsMTx4, does not impair spheroid formation on $Rq_{12}$.

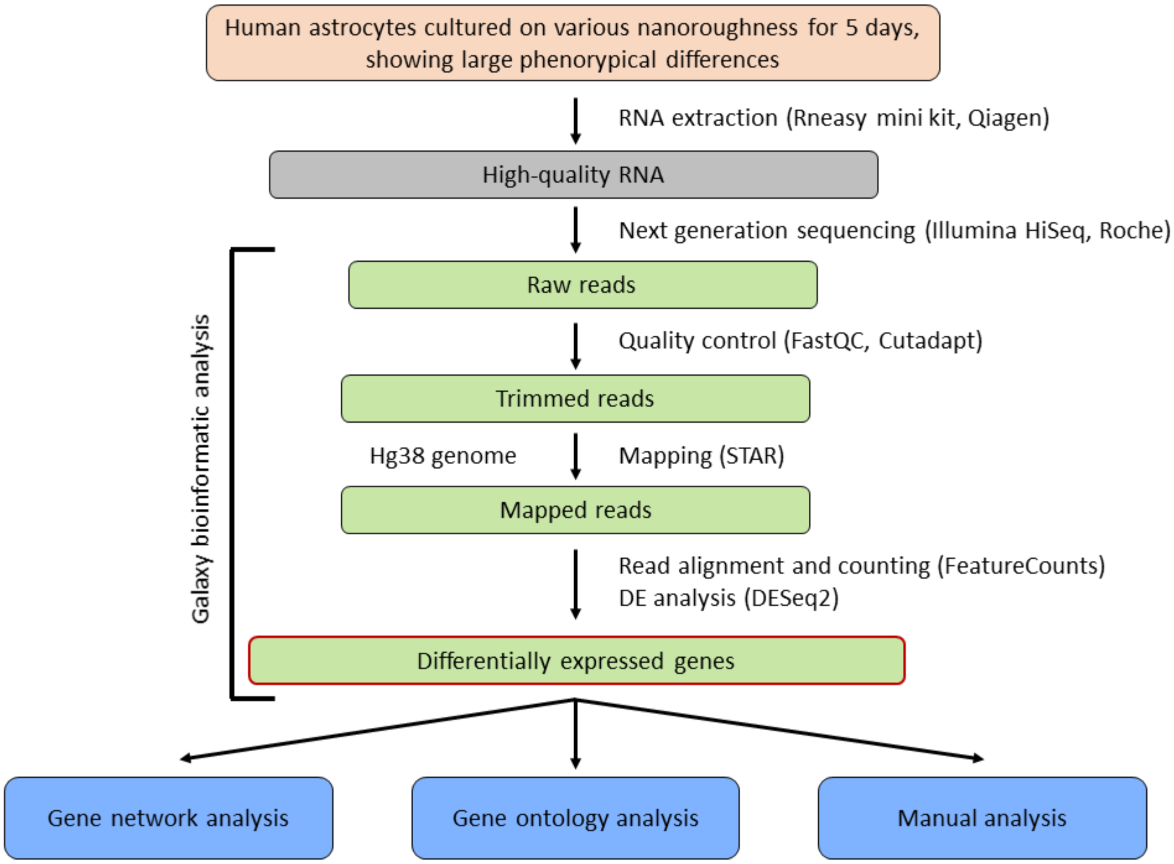


**Fig. S5.** Overview of the RNA-Seq computational analysis pipeline performed using the Galaxy open-web platform.

| | | | | | | | |
|---|---|---|---|---|---|---|---|
| GSKIP | PWAR5 | POLR2M | MTA1 | GULP1 | GDI1 | SULT4A1 | C19orf54 |
| SMARCD3 | HNRNPUL2 | TNFRSF25 | SOX6 | ARFGAP1 | TRO | KRI1 | CAB39 |
| PYROXD2 | PFKL | REEP1 | MAPK11 | SHMT2 | EPHA5 | PLEKHF2 | KMT2A |
| BTG2 | ECE1 | RECQL5 | IFT27 | FLVCR2 | TMCC2 | ATG16L2 | CYLD |
| KCNG1 | USP4 | SPAST | ENO3 | APC2 | ERAP1 | LCNL1 | RHBDF2 |
| MTHFD2L | MIR503HG | TEPSIN | PKIA | PHF23 | CCNG2 | ATF4 | WDR97 |
| DCAF5 | ARFRP1 | DAAM2 | NAB1 | B3GNT2 | NCEH1 | ZNF184 | NAA30 |
| DAZAP1 | MTG1 | RFK | SLC35G2 | HMGB1 | DSG2 | ATG9B | RP11-426C22.4 |
| HS3ST3A1 | PDGFRB | BIN1 | AKAP8L | PLXNA3 | JAG1 | S100A11 | ACER2 |
| KIAA0895L | NTM | BNC1 | TPT1-AS1 | RCN3 | PAN2 | COASY | DMAP1 |
| KBTBD8 | GANAB | PABPN1 | NEURL4 | MMS19 | MMAB | SRP9 | ACIN1 |
| IZUMO4 | PAQR3 | ASPN | ZDHHC21 | CHPT1 | RAPGEFL1 | STAG3L5P | PI4KAP2 |
| TMX4 | CPVL | USP38 | ABHD17B | HECA | ENAH | ELOVL4 | AAED1 |
| GOT1 | PDGFD | CDK18 | TMEM123 | CTD-2547G23.4 | PPTC7 | LSS | ADAMTS13 |
| NR4A2 | FST | AAAS | TP53 | NUFIP2 | RDH5 | INTS3 | MAP1B |
| LRRN1 | BEX2 | ATP6V1F | MT1F | ZDHHC2 | CD58 | A0801 | PLPPR2 |
| LONP1 | YPEL5 | GGH | ZCCHC2 | MAST2 | TMEM165 | ZMYM3 | STAT2 |
| PCOLCE | AGO2 | PTTG1IP | YARS | LZTR1 | NLGN2 | CASP3 | COQ8B |
| UBA1 | UCN2 | LUM | PTPRM | ID3 | SLC25A25-AS1 | UBE2G2 | NHSL1 |
| TSC22D1 | DCT | C12orf49 | TBC1D15 | MSC | VILL | GOLGA7 | GLIPR1 |
| SLC12A4 | NAB2 | CCS | RPL37A | SLC25A37 | CC2D1A | EML2 | ITGA11 |
| SLC25A46 | ACAP3 | FAM200A | SNHG12 | SMARCC2 | PIM1 | TUBE1 | BTBD19 |
| GRK5 | MSANTD3 | C9orf72 | TAF1C | ENPP4 | FAM169A | STK11IP | RP11-66N24.4 |
| SIX5 | GLI1 | ENTPD6 | AGPAT5 | SYNE2 | ACCS | PRKCSH | PLCD3 |
| SEL1L3 | HYPK | NAMPT | ZMIZ2 | VCPIP1 | ADGRL1 | PTMA | WSB2 |
| NEIL1 | ENO2 | IFT172 | FZD3 | FBXL6 | INO80E | UHRF1BP1 | MORC4 |
| LTBP4 | FDFT1 | YAF2 | SOCS5 | SLC9A5 | STK38L | OPLAH | IDUA |
| NCOR2 | NCKAP5L | GPSM1 | PER1 | KIAA0141 | KSR1 | SREK1IP1 | COX11 |
| MIB2 | ITGA6 | MOV10L1 | HDAC10 | DUSP7 | CBLB | CLSTN1 | CIRBP |
| NEFH | USP11 | AC093838.4 | WDR90 | RAB8B | COL6A2 | DPF3 | SGSM2 |
| CCDC57 | LRCH4 | SERPINI1 | LINC00998 | PKD1 | ZC3H12A | SLC11A2 | KLHL17 |
| C6orf15 | CEP164 | NETO1 | SIRT1 | CCDC50 | LUC7L | NBEAL2 | NES |
| SRSF6 | CHTF18 | KLHL15 | LMF2 | FASN | QPCTL | NPHP4 | MICALL2 |
| HDAC7 | NET1 | TYK2 | NEURL1B | MYOM1 | CD248 | PILRB | MMP25-AS1 |
| TRIM2 | PCED1A | ADAMTS15 | PIDD1 | MRI1 | HIGD1A | FOXO3B | TUBGCP6 |
| YWHAQ | DGKQ | MEX3A | ARHGEF25 | NFATC4 | VIM | ZDHHC8 | ADAMTS10 |
| EIF1AX | SLC44A5 | TMEM175 | SBF2-AS1 | TIMP3 | LGALS3BP | LAYN | FAM160B2 |
| TPCN1 | FAM107B | PURA | LARP4 | VPS26A | CACUL1 | FAM83G | COL6A1 |
| STARD13 | SH2D3C | PRICKLE4 | ZMPSTE24 | CYP2U1 | VPS9D1 | TTC31 | FAM193B |
| IFT140 | ANKRD40 | RASA2 | TGFB1I1 | LRRC8C | TTC39C | NCS1 | DDR2 |
| PXDN | CALCOCO2 | OSBPL7 | PYGO1 | SNRNP70 | HSF4 | KLF7 | TRPC4AP |
| RHEB | EPSTI1 | NISCH | NFKBIA | SCOC | SLC36A4 | B3GALNT1 | RNF13 |
| TMEM117 | ARHGAP26 | TNKS1BP1 | TAF15 | RBM33 | EIF5A2 | CPSF1 | HES4 |
| COG1 | ITGB8 | D2HGDH | RNF11 | ATP6V0B | SPRED1 | LIME1 | SRSF5 |
| GTF2IP20 | POFUT2 | FBLN2 | EPAS1 | DPP4 | AC009948.5 | GPR137C | MAGED2 |
| IRS1 | HIPK3 | GOLGA2 | POU2F1 | GGA3 | PSG5 | ANAPC2 | NOTCH3 |
| POMT1 | SREBF2 | RTEL1-TNFRSF6B | ENGASE | CAPZA2 | MFSD10 | MBD6 | PSMB10 |
| FUCA2 | TRUB1 | LPCAT4 | PRPF40B | GPR173 | SPTBN5 | SEC24C | SESN2 |
| IRF3 | ABRACL | PLEKHG5 | DNAJB14 | MINK1 | TMED5 | DENND1B | GLI4 |
| RHBDF1 | APOL1 | RP4-756G23.5 | BACH2 | PIAS3 | SLC12A9 | RP11-802E16.3 | CALML6 |
| SARS | ZBTB41 | C16orf72 | LBH | ARL5B | DENND5B | GGT7 | GALNT16 |
| ATAD3B | CTD-2228K2.7 | RP11-949J7.8 | BMP2K | SOCS6 | FAM118A | KIFC2 | TNS2 |
| PI4KAP1 | ENOPH1 | SLC3A2 | CENPT | ATP1B1 | NXF1 | AMH | AARS |
| KDM4B | MAN2C1 | TRIOBP | COL27A1 | PM20D2 | NSUN5P1 | ZNF460 | NSUN5P2 |
| COL7A1 | SLC5A3 | RHOT2 | CCDC78 | SLC15A4 | ARMC10 | CX3CL1 | FANCB |
| ZNF79 | PCDHB13 | PI4K2B | WNT7B | GRIPAP1 | PPP1R12C | KLHL8 | ARMCX2 |
| NOMO1 | SGSM3 | CDK5RAP3 | ATXN2L | GMCL1 | TTC7A | PPIL2 | TOB1 |
| MTND2P28 | ARSA | STRA6 | MUM1 | PHC3 | TMEM80 | EHD3 | CDK10 |
| SPACA6 | CTD-2270P14.1 | BHLHE40 | ADAMTSL1 | MYO5B | VARS2 | PARP10 | HDAC6 |
| XPC | BGN | BTN2A3P | CCDC126 | POLM | QPCT | PLEC | |
| ASAH2 | SMG1 | EGLN2 | SRRM2 | ZNF692 | ARAP3 | DNAJB9 | |
| FN1 | GNA13 | PJA1 | HNRNPA1 | SPOPL | MST1 | TCIRG1 | |
| GGA1 | TRIP10 | COL16A1 | GIGYF1 | SLC4A3 | MIAT | CDK5R1 | |
| PRPF3 | ZC2HC1A | TP53I11 | GGT5 | COMMD7 | NUP50 | MTCO1P12 | |
| ARID5A | RHBDL1 | MED15 | INSIG1 | SFSWAP | KAT2A | ADCY6 | |

**Fig. S6.** Full list of the 514 DEGs specific to astrocytes grown on $Rq_{12}$.

| | | | | | | | |
|---|---|---|---|---|---|---|---|
| MLC1 | CDC6 | CITED2 | CSTB | CXCL14 | KIAA1217 | TGFB3 | SLC34A2 |
| CXCR4 | STON1 | CEND1 | BOC | PLAU | EFNA1 | SLC2A12 | STC2 |
| GALNT3 | PBX1 | MT1E | PLEKHG1 | CHRDL1 | SLIT3 | CFI | FOXD1 |
| FXYD6 | OLFML2A | HIC1 | LINC01158 | MALT1 | AE000661.37 | ANOS1 | UHRF1 |
| LMNB2 | SLC25A18 | CDON | TMTC1 | GDF6 | WBSCR17 | MT1L | NRG1 |
| CPNE4 | ENTPD2 | EGFR | MT2A | ALK | LCTL | LIMCH1 | ADAMTSL3 |
| STK17A | REPS2 | ARHGEF28 | AOX1 | THBS3 | NTRK3 | LINC00461 | TRDC |
| RP11-342D11.2 | CCND1 | CLSTN3 | PIPOX | DCLK1 | CPED1 | GJB2 | A0800 |
| GMPR | AIF1L | UBE2H | SOX2 | NRP2 | TRIM58 | JAM2 | SELENBP1 |
| PRDM16 | CHST2 | ST5 | SAT1 | PTPRB | LYPD1 | CELSR1 | TRIB2 |
| MEIS2 | PIANP | COL4A6 | CD3EAP | CACNG4 | MOXD1 | PTCHD4 | ADD2 |
| MYO5C | OLFM2 | NHS | PVR | KIAA1549L | MGLL | CADM1 | |
| AFAP1L1 | GPC4 | MARCH4 | PTN | PIK3R3 | PLEK2 | LRRN3 | |

**Fig. S7.** Full list of the 102 DEGs specific to astrocytes grown on $Rq_{32}$.

| | | | | | | | |
|---|---|---|---|---|---|---|---|
| COPRS | SELENOP | ADAMTS6 | SLC1A3 | A0899 | RAB33A | MYO1F | L3MBTL1 |
| C5AR1 | PNOC | DUSP23 | LOX | RP11-274B21.3 | HRASLS5 | RP11-304L19.3 | SLITRK4 |
| COL9A2 | UG0898H09 | FAM111A | TMEM59L | ABHD4 | CASP7 | TMEFF1 | OAF |
| NPC2 | JADE1 | TMEM178B | RP11-158L12.4 | RASL10B | ANKRD9 | GPRIN3 | AVPI1 |
| KLHL30 | AHRR | RPSAP52 | C11orf96 | PLEKHB1 | IL6 | CRTAC1 | LUC7L3 |
| A4GALT | PROSER3 | IQCE | CTSO | LRRC17 | L1CAM | COL10A1 | |
| ADRA2A | LRP1B | MIR31HG | THUMPD3-AS1 | METRN | LEFTY2 | ADAM22 | |
| SPX | KALRN | STAT5B | C8orf33 | TMEM171 | FTH1P20 | SULT1B1 | |
| BAMBI | TPGS1 | CTPS1 | HEXA | EMID1 | CALCRL | RP11-7F17.1 | |
| EBPL | EPHB3 | STARD4 | BAIAP2 | SMOX | TUSC1 | NRSN1 | |

**Fig. S8.** Full list of the 75 DEGs specific to astrocytes grown on $Rq_{16}$.

| | | | | | | | |
|---|---|---|---|---|---|---|---|
| PNMA1 | ZMIZ1-AS1 | NXN | EPS8L2 | SYNPO2 | CAMKK1 | MIR4697HG | ERCC6 |
| CD44 | LINC01605 | ZNF365 | RPL35A | EPHA2 | SLC9A7 | PHLPP1 | GCNT4 |
| C10orf90 | HRH2 | RP11-543P15.1 | SNHG19 | SYNGR3 | TUFT1 | SLC4A10 | KCNC4 |
| ITGA5 | PRSS22 | FAHD2B | DCXR | SMAD3 | ANO1 | VDR | RPL23 |
| SALL1 | GPR158 | FAM184B | LURAP1L | SLC2A13 | GSN | CPNE8 | AKAP5 |
| ADAM12 | PACSIN3 | RPS6 | ILDR2 | FBN2 | EGR2 | WDR81 | CYTH3 |
| RPS18 | C1orf198 | PKN3 | SRA1 | RFTN1 | SH3KBP1 | RPL26 | RGS6 |
| LINC00520 | LXN | PFKFB2 | SSPN | ATP8B2 | ANK2 | DIRAS3 | RPL13A |
| KCNK3 | THY1 | PNMA2 | IGSF9B | MURC | PLA2G4A | RPS2P7 | RPS7 |
| KCNK6 | TBC1D8 | ATPIF1 | LCP1 | SLC25A1 | SELENOW | KCNT2 | PLEKHS1 |
| C4orf3 | SHH | ISLR | PCDHGB6 | BDNF | ITGB6 | AFF2 | RGS10 |
| LRRN2 | CNTNAP3B | POSTN | NRXN3 | HBEGF | PPP1R3B | RACK1 | QSOX1 |
| BET1L | C5orf46 | SPRY1 | AJUBA | TFCP2L1 | ZBTB16 | PRG4 | FRZB |
| SORL1 | SUSD6 | OSGIN1 | ADM | YEATS2 | MPI | SLC36A1 | COL6A3 |
| RPL7A | AC156455.1 | PTPRF | SNHG6 | MGAT1 | RNF112 | SDCBP | PLEKHA4 |
| FRY | GLIPR2 | CASC15 | RPL11 | ULK1 | ADAMTS4 | IGF2 | KCTD16 |
| RP1-140K8.5 | NEFL | NDRG1 | SNX8 | ZNF175 | STK32A | RPL21 | KIF20A |
| CXADR | C6orf48 | EEF1D | RPL8 | PTHLH | MYH10 | CLIC3 | MIF |
| DKK2 | RNF150 | RP11-9G1.3 | RGS17 | C8orf4 | C14orf159 | HOMER2 | WDR86 |
| EPS8 | CLIP2 | KLHL38 | MBNL1-AS1 | MAP3K14 | STXBP5 | PRKG1 | CHRNA9 |
| LINC00460 | RPL3P4 | RPS3A | RPL23A | NOVA2 | IFNGR2 | FGF7 | HIVEP3 |
| VKORC1 | MARCH3 | LRP8 | WIPF1 | COPZ2 | SLC44A1 | FMNL3 | BMPR1B |
| TEK | CTSK | RPL34 | ST6GAL1 | GALNT5 | OLAH | RPL5 | SHC4 |
| DGUOK | CD93 | HS6ST1 | RPS25 | RPS4X | GPRC5A | ZDHHC9 | XYLB |
| NECTIN2 | RASSF9 | AKAP12 | EFR3A | NIPSNAP1 | VWF | MTURN | KITLG |
| RHOC | CDK4 | TTL | OST4 | RPS3 | EBF1 | TSPO | IGFBP4 |
| RPL21P16 | CALB2 | RPL18A | BAALC | RPS27A | ITM2C | ARRDC3 | FMNL2 |
| CORO2A | CPA4 | LIPG | RPS2P5 | HOXB3 | SELENOM | TCAF2 | MAMDC2 |
| GATA6 | GNG2 | SP140 | GALNT1 | IGFBP7 | AMDHD2 | FAM46A | REXO2 |
| DSTN | PDE3A | APOLD1 | PPARGC1B | RP11-513I15.6 | TNS1 | DOCK10 | TM4SF19 |
| RPL18AP3 | TRHDE | APOD | FAM13B | ADGRL2 | EIF3F | IGDCC4 | RP11-253M7.1 |
| DNM1 | MFAP3L | ARHGAP29 | RPL38 | SLC47A1 | DOK6 | RPS12 | C4orf26 |
| RPLP1 | SLC7A11 | OSBPL1A | RPS16 | PAPPA | G0S2 | C19orf48 | SPINT2 |
| TOX3 | SLC37A1 | ACVR1 | ZBTB46 | RPS13 | EEF1B2 | IAH1 | FDPS |
| AC004988.1 | MAPK8IP2 | COMP | RPL14 | ZP1 | SEMA3B | SOX8 | SNAP91 |
| RGL1 | RPS24 | RPL3 | HEPACAM | LINC00862 | MRPL17 | AEBP1 | NANS |
| OXTR | RPL27 | LOXL1-AS1 | LRRC75A-AS1 | STOX2 | RPL6 | PRR5L | CDH5 |
| KIAA0355 | RPL24 | PTPRG | ARSJ | RPS8 | AK2 | SLC9A1 | MROH1 |
| ZNF395 | GBX2 | TXNRD1 | TMEM256 | HHIP-AS1 | PLCE1-AS1 | WNT2B | NPC1 |
| PRDX4 | PKP2 | LACC1 | ALPL | KIAA0196 | NIPAL2 | RPL18 | GRAMD3 |
| GLIS2 | NME4 | RPL9 | RUBCNL | EGFL7 | ARRDC4 | HOXB6 | ECSCR |
| RPL32 | EFR3B | GRIK3 | ACKR3 | NEK7 | CPPED1 | SNHG8 | APP |
| SPG20 | UCP2 | TNIP1 | LTA4H | EMP1 | MR1 | ST6GAL2 | CSPG4P13 |
| VASH1 | FER1L6 | CCDC85C | FNBP1 | RRAS2 | TXNDC16 | RPS19 | APBA1 |
| FKBP4 | ARHGAP20 | RAC3 | CKAP4 | LTBP2 | ARHGAP23 | RPL27A | |
| NDNF | INPP5F | PCDH17 | COBLL1 | SYNGR2 | BMPER | SLC41A2 | |
| LOXL1 | VASH2 | HIP1R | RPL10A | RCOR2 | CCND3 | TGFA | |
| PRRX1 | GGT1 | PLEKHA2 | RCBTB1 | KDM1A | FLJ22447 | ZFYVE26 | |
| IFI16 | ZCCHC6 | RP11-96H19.1 | ADGRB2 | ILK | HMGB2 | GDF5 | |
| NUAK1 | NECTIN1 | NDUFB11 | FOXQ1 | CARD6 | MGARP | GSTM3 | |
| LAMB3 | RPLP0 | ADAMTS3 | DOCK2 | FABP3 | RPL17P50 | NFATC2 | |

**Fig. S9.** Full list of the 75 DEGs specific to astrocytes grown on $Rq_{24}$.

| | | | | | | | |
|---|---|---|---|---|---|---|---|
| PKNOX2 | ALDH1L2 | RP4-555D20.2 | NID2 | HOPX | DTX1 | PML | MEG3 |
| KLHL13 | GPRC5C | BRICD5 | RAB22A | MAP4 | FBXO33 | LAMB1 | NEDD9 |
| APOL2 | RP11-574K11.24 | CRTAP | MZF1 | RPL36A | CCNL2 | STARD10 | NRBP2 |
| A0802 | IL18BP | GTPBP2 | THRA | CDH13 | TSPYL2 | PABPC1L | ECHDC2 |
| TRIB3 | RBM6 | SLC26A6 | CAMTA2 | TMEM86A | ELN | NAT9 | HEMK1 |
| SP140L | NEK6 | ITGA10 | DGKA | MXRA8 | SAMD9L | EIF4A1 | TBC1D2 |
| CERK | PIGL | PNISR | GALT | ATF3 | EPRS | CCDC18-AS1 | MALAT1 |
| SPPL2B | GABBR1 | IQGAP3 | ALDH1A3 | APOL6 | C11orf95 | MEG9 | LHX2 |
| SLC26A10 | CLTCL1 | ZNF767P | HSPA7 | PAQR6 | MEGF9 | DOK3 | PNN |
| C15orf52 | VMP1 | COL5A3 | MAMDC4 | DNAJA4 | LENG8 | HSPA5 | PRSS53 |
| ANKRD6 | AFG3L1P | HNRNPH1 | CARMN | DACT3 | SHC3 | CSAD | NDFIP1 |
| APOE | PHYKPL | FNDC4 | LINC01583 | UBA7 | BEX4 | BTN3A3 | FLNA |
| PLEKHG4 | MEMO1 | PCDHB14 | CEACAM19 | LINC00511 | LAMA5 | ANPEP | FCHSD1 |
| BTN3A1 | NFKBIZ | MIR34A | FAM210B | SEC31B | TLR3 | TTLL3 | RP4-800G7.2 |
| ID1 | XAF1 | APBB3 | MAPK8IP3 | GPI | PTGDR2 | RP11-159D12.2 | GOLGA8B |
| FUS | EME2 | GNAI1 | MICAL1 | RNF145 | AC005154.6 | SUGP2 | DCAF8 |
| RP11-66N24.3 | RP11-69E11.4 | PIF1 | SMG1P7 | ARHGAP33 | MTATP6P1 | TUBG2 | |
| CYP1A1 | PLPP2 | IARS | LCAT | PLEKHG2 | TRPM4 | CBLN3 | |
| LTBP1 | MIR222HG | SLC6A8 | ACTN1 | BAG1 | ITGA4 | SERPINE1 | |
| ZBTB20 | VEGFA | RBM5 | PSME2 | DMPK | THAP2 | EWSR1 | |
| C15orf59 | PSEN2 | GSDMB | GNB3 | NEAT1 | ODF2 | NCALD | |

**Fig. S10.** Full list of the common DEGs between astrocytes grown on $Rq_{12}$ and $Rq_{16}$.

| | | | | | | | |
|---|---|---|---|---|---|---|---|
| COL4A5 | SPP1 | SLC6A1 | LDLRAD4 | PRUNE2 | DGKI | DKK1 | ACTA2 |
| A2M | NAV2 | SLC37A2 | TIPARP | GREM1 | SLC16A6 | VCAM1 | SYNPO |
| SLC6A15 | GABRE | FZD8 | CFH | DHRS3 | FOXRED2 | TNFAIP2 | AHNAK2 |
| SLCO2A1 | LAMA1 | LOXL4 | KIAA1755 | CCL2 | SORBS1 | CMB9-55A18.1 | SH2D5 |
| RARRES3 | E2F7 | JAKMIP2-AS1 | ADGRL4 | HMGA2 | BRI3 | NRCAM | ITPR3 |
| ELFN1 | CA12 | CYP1B1 | STC1 | SULT1E1 | FNDC5 | NPTX1 | |
| RASSF4 | GALNT15 | PNP | FER1L4 | TMEM158 | RALA | CDCP1 | |
| TNFAIP8L3 | HMOX1 | GPNMB | SPHK1 | TNC | NKD2 | SHISA2 | |
| EPHA7 | SIGLEC15 | VAMP1 | SEMA7A | INPP4B | PCDH10 | ENC1 | |

**Fig. S11.** Full list of the common DEGs between astrocytes grown on $Rq_{12}$, $Rq_{16}$, $Rq_{24}$ and $Rq_{32}$.

| | | | | | | | |
|---|---|---|---|---|---|---|---|
| FSTL1 | CYSTM1 | WEE1 | HTRA1 | TMC7 | SLC1A5 | GLMP | THBS2 |
| CLMP | SORBS2 | CD109 | ID2 | ZNF697 | ITGA8 | CHAC1 | PRSS23 |
| CREB3L1 | PCDHGC5 | ADAMTS12 | ROBO3 | SLC17A5 | RUNX1 | ADGRA2 | ASNS |
| KCNMA1 | LMO4 | PLXNA4 | PTPRN | ITGB5 | FADS2 | UBE2D1 | CXCL1 |
| CNTNAP1 | ARHGEF16 | GLCCI1 | INHBE | ZNF503 | MID1 | ZCCHC14 | ATP6V1G1 |
| EDN1 | SNN | ADGRG1 | MAP3K4 | KCNS3 | CXCL8 | ACSS2 | LOXL3 |
| IRAK1 | PROX1 | MMP11 | HEY1 | ARL8B | HDAC9 | RP5-1172A22.1 | FAT3 |
| GAS5 | RP11-268J15.5 | B3GALT2 | GPAM | MEDAG | TOM1L2 | STX3 | SPON2 |
| FAM134B | TNFAIP6 | ANXA2 | SRPX | CHST3 | DYRK2 | RGS2 | ERF |
| MMP7 | SNAI1 | PLPP4 | MAFB | VGLL4 | TARS | NXPH4 | S100A10 |
| RDH10 | SLC17A9 | STK10 | RP11-863P13.3 | TRIM62 | ABCA8 | PDPN | RUSC2 |
| GYS1 | VSTM4 | SHC2 | SGCD | XYLT1 | NDUFA4L2 | COL3A1 | HSPH1 |
| ERG | CSRP1 | MDGA1 | FILIP1L | MIR210HG | BNIP3 | KDR | CYP26B1 |
| SOD2 | CTSB | ADARB1 | ENO1 | SLC19A2 | LEF1 | PRKAR2B | IGFL3 |
| DIRAS1 | EHBP1L1 | CNN2 | P4HA1 | ARTN | MTHFD2 | DNM3 | CREBRF |
| HCN1 | PGAM1 | LMO7 | COL4A1 | BEAN1 | ANXA6 | OGT | FAM162A |
| PDLIM2 | EEPD1 | COL4A2 | OSBP2 | CERS4 | RAB27B | TUBA4A | CDH23 |
| FIBCD1 | CACNA1H | PPP1R13L | PLPP3 | ANKZF1 | LYST | RNF207 | TGFBR2 |
| FADS1 | UNC5B-AS1 | C1S | LAMB2 | MCTP1 | RAMP1 | FAM13C | IL33 |
| GPR153 | LDLR | BMF | SLC39A10 | MEF2C | PLEKHG3 | P4HA2 | PDK1 |
| LIMS2 | DDI2 | FLT1 | KIAA1211 | PSPH | FKBP10 | MOV10 | FLG |
| ACSM3 | SIX2 | DYRK3 | MYL9 | PGK1 | SMAD7 | STXBP2 | C6orf132 |
| GLT8D2 | COL5A2 | LINC01050 | LAMP2 | PLOD1 | VLDLR | CBFB | MTSS1 |
| IMPDH2 | PGM1 | EBI3 | SFTA1P | ESM1 | FHL2 | IFITM3 | ANGPTL2 |
| TENM4 | RCAN1 | LRRC32 | ADAM17 | B3GNT5 | WARS | RBCK1 | ARHGEF17 |
| MCUB | NTN4 | RPL41P5 | TPM2 | KIF26B | GOLGA8A | PCDH1 | APCDD1L-AS1 |
| COL1A1 | HSPA2 | CD276 | MGAT3 | TSPAN11 | DES | MN1 | ENPP1 |
| AP1S2 | KRT8 | F13A1 | MBP | RENBP | RP3-428L16.2 | ATP6V1B2 | UNC13C |
| ALDOC | RAB20 | ANO4 | CCDC71L | MMP13 | TMEM35A | CD24 | HIPK2 |
| AC112721.1 | PHGDH | DUSP3 | RASSF8 | FAM109A | LMO2 | ARHGAP9 | SLC6A9 |
| TSPAN13 | EHD2 | FLCN | MAF | PKP1 | LDHA | PSAP | SYT14 |
| DYSF | CBS | PTGER2 | DOK5 | HOMER3 | EDNRB | PTGDS | NOV |
| NKD1 | BEGAIN | HS3ST3B1 | DMD | KLF11 | SCD | HERPUD1 | PPP1R3C |
| NAT8L | SULF2 | CTNS | RP5-1054A22.4 | IRF1 | MAN1C1 | BCL2L11 | GPM6B |
| DUSP4 | LAMC3 | FAM102A | NUAK2 | FPR3 | HSPA6 | SCPEP1 | RRAGD |
| ALDH18A1 | MGP | CCDC3 | MANBA | SOX4 | GM2A | SERPINE2 | SOX11 |
| TPI1 | MVD | FAM171B | HAX1 | RNF41 | CRISPLD1 | METTL9 | SIX1 |
| KIAA1211L | SYT7 | FNIP2 | CHRM3 | C8orf58 | CDC42EP2 | FAM198B | ITGB3 |
| PLIN2 | KCNN3 | PDLIM1 | CPE | TRIM22 | H19 | SDPR | MARS |
| FNDC1 | RRAGC | PYGB | SLC24A3 | TGM2 | MCHR1 | MYRF | P4HB |
| HS2ST1 | TPPP3 | SH3PXD2B | ST3GAL5 | CLDN4 | UACA | LOXL2 | TLL1 |
| CHMP1B | ZFAS1 | TCEAL7 | ARHGAP24 | PDLIM7 | HEXB | ADM2 | TMEM132B |
| SYT12 | DENND2A | SCG2 | SORBS3 | SLC44A2 | TPM1 | CADM3 | PPP1R14C |
| AC133644.2 | GPRIN2 | MMP14 | CPT1A | SQSTM1 | DIEXF | SLC7A5 | IL11 |
| PGGHG | GPR1 | ITGBL1 | PFKFB4 | TMEM127 | CRLF1 | UBL3 | GPRC5B |
| P4HA3 | MXRA5 | CPXM1 | SREBF1 | UAP1L1 | FLNB | AK4 | TCF7 |
| ABHD2 | TRPC4 | MYLK | DCN | APCDD1L | CTH | GUCY1A2 | AKR1C3 |
| C14orf132 | NR1D2 | MAPKAPK3 | DLK1 | ANKRD10 | CLEC18B | TPP1 | SPARC |
| PCK2 | GAP43 | B4GALNT3 | PLVAP | ST3GAL1 | PYCR1 | ENPP5 | VIPR1 |
| CHST11 | PROS1 | ZFP36L1 | MIR24-2 | CTIF | COL4A3BP | CLCN5 | KCTD12 |
| MMP15 | TINAGL1 | SNX9 | COL11A1 | AC007255.8 | LRRC15 | AP1G2 | CTSH |
| ASS1 | ITGA2 | FAM134A | GPR68 | XIRP1 | OSTM1 | CKLF | COL5A1 |
| CTD-2269F5.1 | VAT1 | SIRPA | FOXF2 | LGALS1 | RRBP1 | SDC1 | GPT2 |
| ADAMTS16 | GRIN2A | IGFBP6 | AC112721.2 | GNPDA1 | CD9 | SLC1A4 | PTGS1 |
| EDNRA | FAM213A | CSMD2 | COL8A2 | DRAXIN | CTSD | HPCAL1 | PPFIA4 |
| CDC42EP3 | KANK4 | SAMD11 | TIMP4 | STEAP3 | ADCY8 | SPG21 | CNIH3 |
| LRRC8A | MYC | C9orf172 | HSPB7 | CLEC2B | CDKL2 | PSAT1 | TMEM119 |
| GRIK2 | PHYH | LRRTM2 | GAPDH | CRYAB | CAMK2N1 | CREG1 | RP11-448G15.3 |
| PHLDA1 | TIMP1 | PDGFA | PMP22 | FNIP1 | GRIA3 | CCDC80 | PYGM |
| AGT | POU3F2 | GBP1 | B4GALT5 | ABC7-42404400C2 | LARP6 | SPOCK2 | HKDC1 |
| FGF1 | PCSK9 | OPCML | BPI | ATP13A3 | FBN1 | CLU | DNER |
| NFIL3 | RDH11 | SH3D21 | MAGED1 | SLCO3A1 | C7 | CXCL12 | MFHAS1 |
| TM6SF1 | AMOTL2 | SCARA3 | ETV1 | DUSP2 | TYRO3 | NTN1 | POPDC3 |
| CHI3L1 | KIF5C | C5AR2 | PEAR1 | MEF2A | RAB31 | TMEM173 | GEM |
| CMTM3 | DUSP10 | ARMCX1 | SYNM | CAMK2D | SOAT1 | RAB29 | KCNJ15 |
| RP11-400N13.3 | GPM6A | ACSL1 | MYO1D | COL21A1 | SPAG4 | SLC20A1 | STK38 |
| NEU1 | IBSP | TNFRSF21 | HSD17B14 | HMCN1 | ALDOA | KHDRBS3 | FBXO32 |
| SLC9A3R1 | GCLC | BMP1 | ACTG1 | STAT4 | HEG1 | UBE2E1 | KCNN4 |
| CARS | IGFBP5 | TMEM38B | KDSR | ZMYND8 | TXNIP | COL14A1 | PLXDC1 |
| MGST1 | COL8A1 | TBX3 | FKBP11 | RASGRP2 | SLC7A8 | PRR15 | ADGRA3 |
| ASAH1 | EPHB1 | RASL11B | LPCAT1 | ARHGEF2 | M6PR | SNX30 | RNF128 |
| NADSYN1 | MEGF6 | MTHFD1L | COL1A2 | STRIP2 | ZNF521 | CAV1 | APOL3 |
| FHL1 | NRROS | KRT18 | CYR61 | ITGA1 | CTGF | RAB3IL1 | HEYL |
| LPIN3 | LSAMP | TMEM200C | DTNA | PPARG | CMAHP | SCARB2 | CP |
| SERTAD2 | PTGER4 | SYT11 | IL1R1 | RNF144A | PSME1 | CTSL | |
| NPTXR | GARS | KLHL24 | RAB7A | SPOCK1 | MBOAT2 | PGPEP1 | |

**Fig. S12.** Full list of the common DEGs between astrocytes grown on $Rq_{12}$, $Rq_{16}$ and $Rq_{24}$.

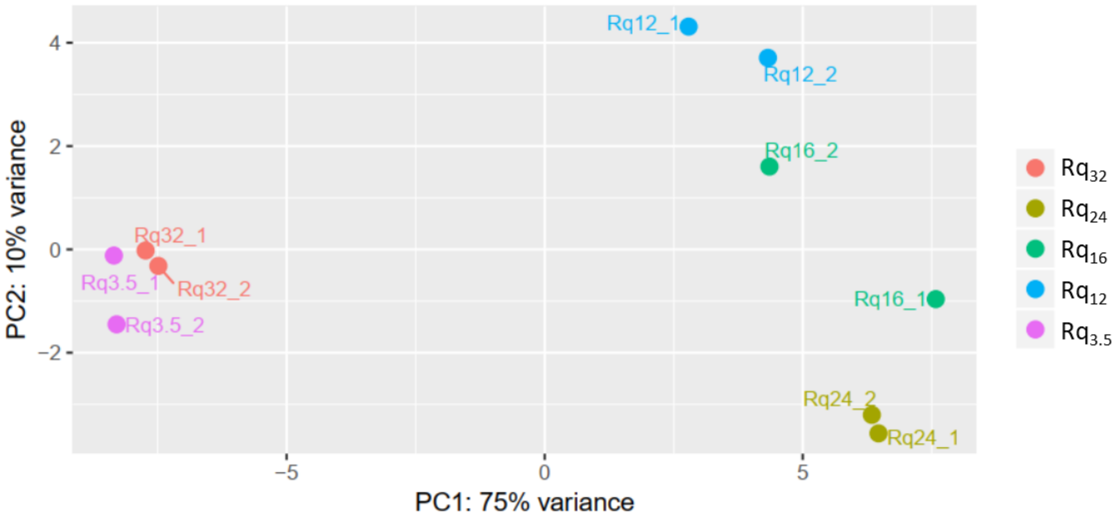


**Fig. S13.** PCA plot from the DESeq2 (3.5.1) analysis. The x-axis PC1 represents the effect of nanoroughness on astrocytes while the y-axis PC2 is a theoretical side factor. The PCA plot highlights the similarity between the gene expression profiles of astrocytes grown on both extremes of the nanoroughness range, $Rq_{3.5}$ and $Rq_{32}$, differing significantly from $Rq_{12}$, $Rq_{16}$ and $Rq_{24}$. Within the latter three, $Rq_{12}$ shows a unique trend.

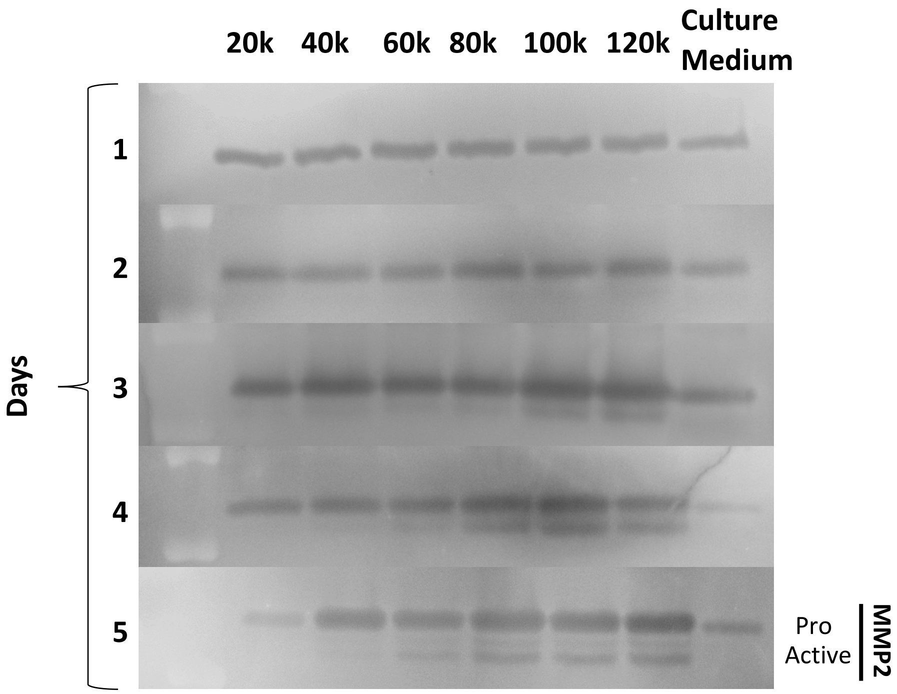


**Fig. S14.** Time course quantification of MMP release in the culture medium by different numbers of astrocytes seeded on $Rq_{12}$ over 5 days by zymography. The top band at 72 kDa represents pro-MMP2 while the lower band shows active-MMP2. MMP9 is not shown as no differences were observed.

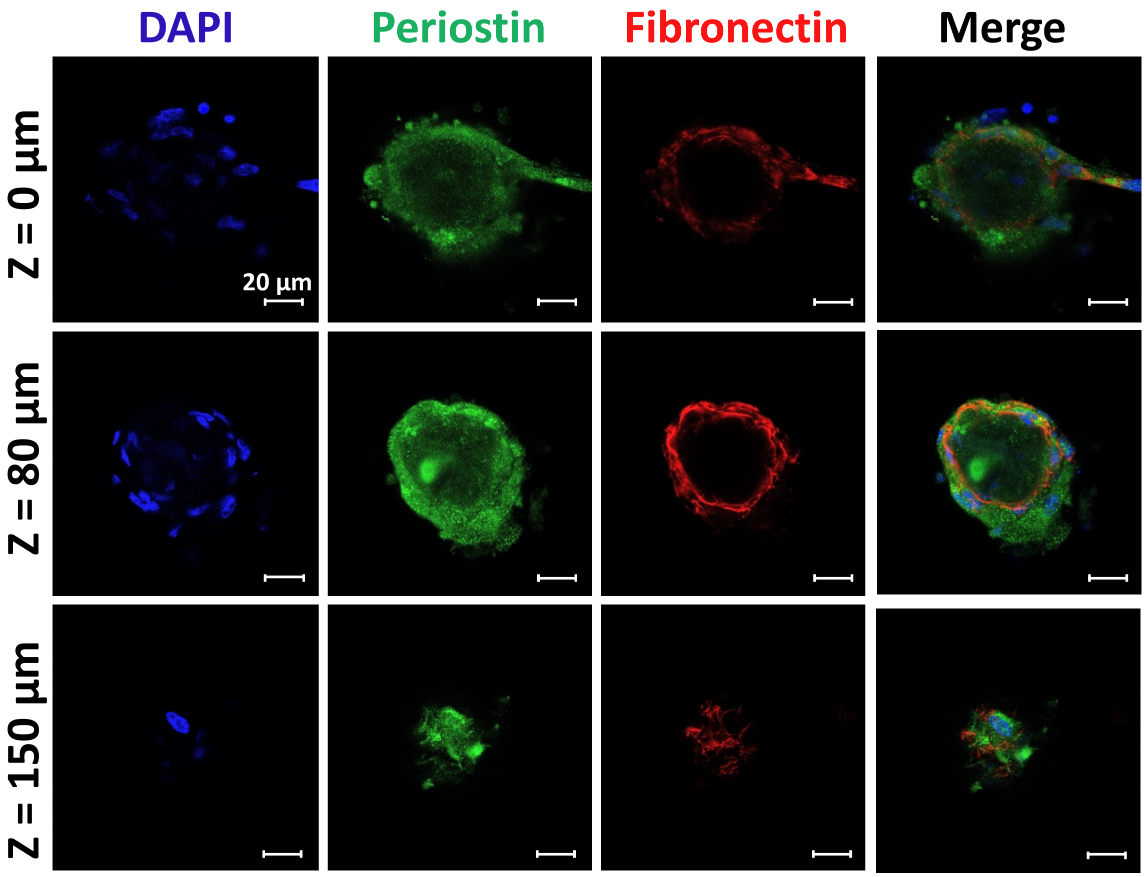


**Fig. S15.** Confocal imaging of an astrocyte spheroid on Rq12, stained for periostin and fibronectin.

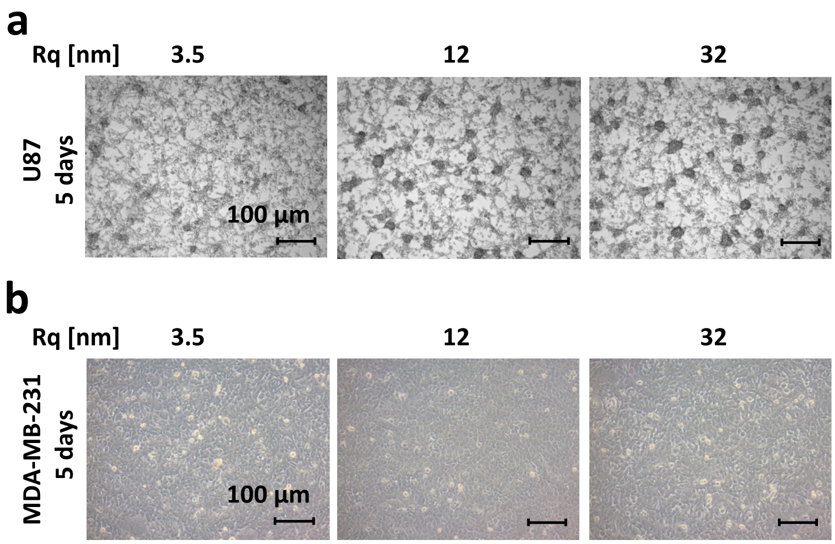


**Fig. S16.** Culture of cancer cells on nanoroughness **(a)** U87 cells do not exhibit phenotypical differences between the different nanorough substrates **(b)** MDA-MB-231 cells form a cellular monolayer regardless of the underlying nanoroughness.

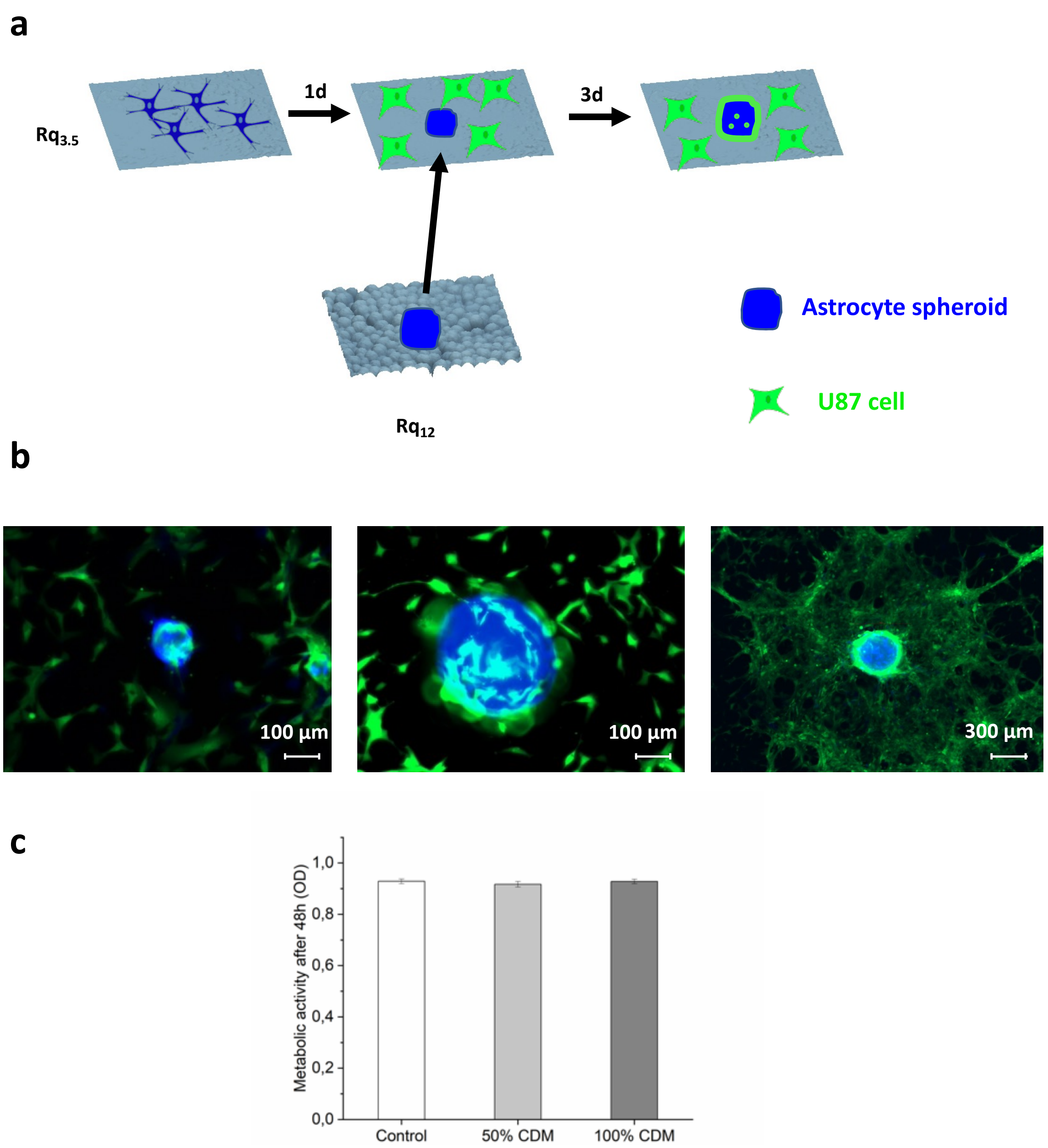


**Fig. S17.** Further investigation of the crosstalk between astrocytes and U87 cells. **(a)** Schematical representation of the experimental design. Astrocyte spheroid formed on Rq12 were trypsinized and added to U87 cultures on smooth $Rq_{3.5}$ substrate. **(b)** U87 cancer cells after 3 days of co-culture showed strong spatial association with the astrocyte spheroids. **(c)** Assessment of metabolic activity of U87 cells after 2 days growth in medium conditioned by astrocytes grown on $Rq_{12}$.

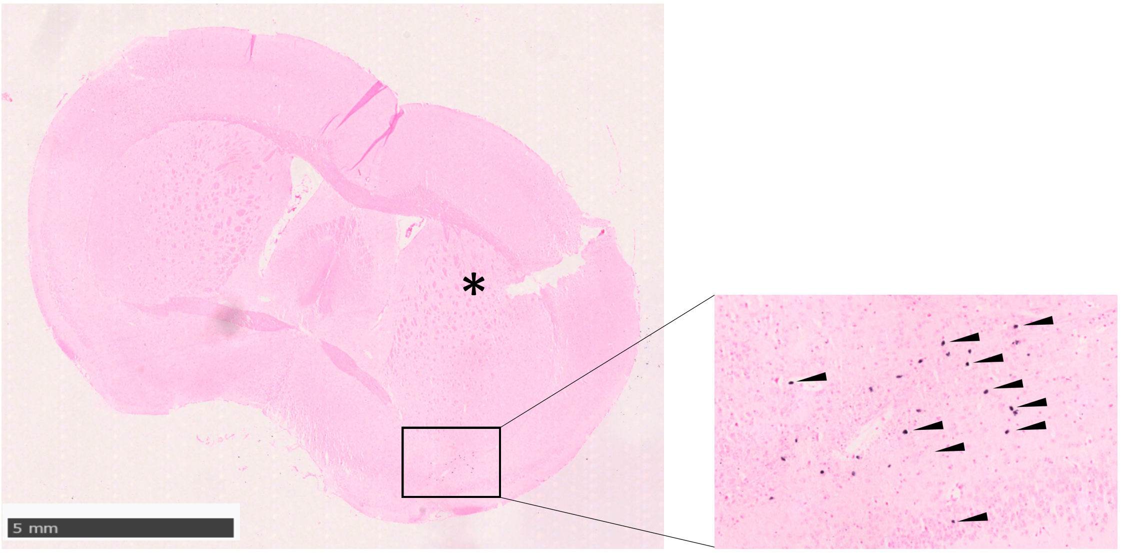


**Fig. S18.** Coronal section of a representative mouse brain injected with dissociated spheroids of astrocytes grown on $Rq_{12}$. Human astrocytes (blue nuclei) were identified far from the injection site (indicated by the star) 12 months after injection by in situ hybridization for the Alu human sequence. The arrowheads in the higher magnification of the outlined area point toward some Alu-labelled human nuclei.
